# Flower patterns improve foraging efficiency in bumblebees independent of nectary guidance

**DOI:** 10.1101/2022.06.20.496816

**Authors:** Robin Richter, Alexander Dietz, James Foster, Johannes Spaethe, Anna Stöckl

**Affiliations:** Behavioral Physiology and Sociobiology (Zoology II), University of Würzburg, Biozentrum am Hubland, 97074 Würzburg, Germany

**Keywords:** flower pattern, nectar guide, pollination, foraging, insect

## Abstract

Colourful patterns on flowers are thought to benefit both pollinators and the plants they visit, by increasing the plants’ pollination success via an improved foraging efficiency of its pollinators. This increased efficiency is thought to result from a guidance effect of the flower patterns, correspondingly termed ‘nectar guides’, which indicate the position of the nectary to visiting pollinators. While it is well established that flower patterns play an important role in flower choice, the mechanisms underlying their function for flower-visiting insects remain poorly understood. In this study, we quantified the contributions of patterns to all phases of flower interaction in the buff-tailed bumblebee (*Bombus terrestris*). We analysed their flight paths, as well as landing positions and walking tracks on artificial flowers with different pattern types. We reveal that flower patterns improved the overall foraging efficiency of the bees by up to 30%, by guiding their approach flight, landing positions, and departure decisions. Surprisingly, these effects were not related to nectary guidance. Since we conducted the experiments with experienced foragers, which represent the majority of insect pollinators active in nature, the newly described nectary-independent guidance effects of flower patterns are of fundamental importance to plant-pollinator interactions under natural conditions.

## Introduction

We have all experienced the fascinating range of colourful patterns that flowers display on our windowsills and in our gardens – though while merely pleasing to us, they can be of great importance to animals that visit flowers for their daily food supply (Dafni & Giurfa 1999, de Ibarra et al. 2015, van der Kooi et al. 2019). Already in the late 18^th^ century, Sprengel proposed that these “sap marks” (*Saftmale*) may guide pollinators to the nectary (Sprengel 1793). The assumption that such *nectar guides* mutually benefit pollinators and the plants they visit is supported by the presence of these patterns within numerous plant families, which possess colour and contrast ranges that are readily detected by insects (Chittka et al. 1994, Dafni & Kevan 1996, Dafni et al. 1997, De Ibarra & Vorobyev 2009, Penny 1983). On the other hand, pollinators of various insect groups show innate preferences for naturalistic flower patterns, in particular for radial patterns and central elements (Dafni et al. 1997, Hansen et al. 2011, Heuschen et al. 2005, Johnson & Dafni 1998, Kelber 2002, Lawson et al. 2017, Lunau et al. 2009).

Pollinators are thought to benefit from flower patterns via two main mechanisms: flower patterns help them to memorise and select nectar-rich flowers, in addition to other visual and sensory cues (de Ibarra et al. 2015, Latty & Trueblood 2020, Orbán & Plowright 2014, Stöckl & Kelber 2019), and they can improve the efficiency of flower visits, by reducing handling time. Distinctly shorter handling times with patterned artificial flowers, as compared to uniform ones, have been observed in hymenopteran, dipteran and lepidopteran pollinators (Dinkel & Lunau 2001, Goodale et al. 2014, Goyret 2010, Goyret & Kelber 2012, Johnson & Dafni 1998, Lawson et al. 2017, Leonard & Papaj 2011, Waser & Price 1985). For the plants, both an increased attractiveness, and higher visitation rates (due to shorter handling times) result in an increased chance of pollen dispersal (Nepi et al. 2018, Valenta et al. 2016).

While the role of patterns for both innate and acquired flower selection are well established (de Ibarra et al. 2015, Latty & Trueblood 2020, Orbán & Plowright 2014, Stöckl & Kelber 2019), the mechanisms by which patterns increase foraging efficiency, measured as the ultimate reduction in time spent moving between flowers (Waser & Price 1985) or on the flower preceding nectary discovery (Goyret 2010, Goyret & Kelber 2012, Lawson et al. 2017, Leonard & Papaj 2011), still remain largely elusive. To understand these effects, it is important to subdivide the flower visit into its component stages: the approach flight, the choice of landing position and control of landing, the movement across the flower towards the nectary, and the decision for take-off after nectar uptake (Fig. 1A). Very little is known about the role of flower patterns in the stages before and during landing. Previous evidence suggests that honeybees and bumblebees only recognise patterns in the immediate vicinity of flowers (de Ibarra et al. 2002), and do not use them to control their approach flight over longer distances (Free 1970, Manning 1956). There are indications that bees land more centrally on a flower with a pattern in its centre (Free 1970, Manning 1956) – though this was tested with flowers distinctly larger than those visited by bees in nature. Thus, to reveal if and how patterns guide the first two stages of flower interaction - approach and landing, landing behaviour must be tested with a variety of flower patterns on naturally sized flowers.

**Fig. 1.**
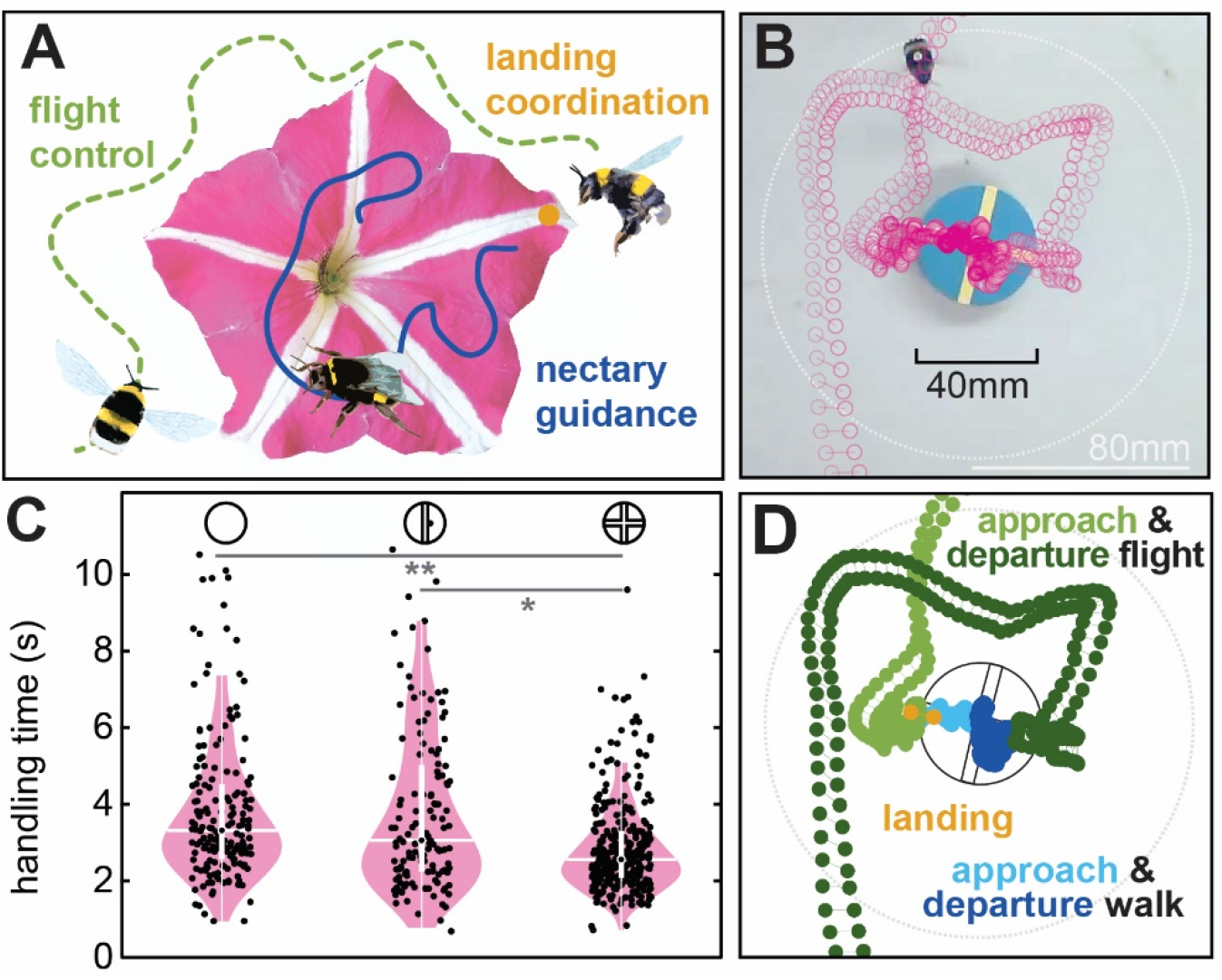
The functional significance of flower patterns for bumblebee foragers. **A** Flower patterns provide important visual cues for several stages of flower interaction: for orientation and flight control during the approach flight, for landing control, and as a visual guide to the nectary. **B** We investigated the functional role of flower patterns in these different stages by filming the entire interaction of bumblebee foragers with artificial flowers, which were mounted horizontally 10 mm off the ground, and presented flower patterns of different shapes and colours (see Fig. S1). The head and thorax position of each individual bumblebee were tracked at 60 fps. **C** Total flower handling time of bumblebee foragers (from approaching the flower at a perimeter of 80mm, shown in **B**, to take-off, excluding time during nectar uptake) with three different flower patterns: *no*, *line* and *cross* pattern. The asterisks depict the statistical results of a generalised linear mixed-effects model (GLMM, see methods, Table S1). Only significant results of the pairwise comparisons of the *pattern-type* and *foraging-time* factors are shown. **D** To investigate which stages of flower interaction contributed to pattern-dependence of flower handling time, and what their underlying mechanisms were, we analysed different stages of flower interaction: the approach (and departure) flight, the landing position and orientation, and the walking paths before and after nectar uptake.

In contrast to the approach and landing phase, there are clear indications that flower patterns guide the third stage of flower interaction: the pollinator’s movement across the flower to the nectary. Bumblebees, which typically walk to the nectary after landing, required less time to find the nectary on realistically sized flowers when patterns were present (Leonard & Papaj 2011). Similarly, hovering hawkmoths require shorter exploration times to guide their proboscis to the nectary when radial patterns are present (Goyret 2010, Goyret & Kelber 2012). However, in bumblebees, this effect only occurred when the bees were trained on patterned flowers and then presented with uniform ones, not vice versa. Furthermore, the effect decreased over a limited number of consecutive flower visits (Leonard & Papaj 2011), and thus strongly depended on experience. Considering that honeybee and bumblebee foragers perform hundreds if not thousands of flower visits per day (Abrol 2012, Pasquaretta et al. 2019), it remains an open question what, if any, advantage floral guides convey to experienced foragers, which perform the majority of flower visits. Interestingly, flower patterns also appear to play a role in the last stage of flower interaction, after nectar collection, in reducing the time naïve bumblebee foragers spent exploring artificial flowers before take-off (Leonard & Papaj 2011). Given that considerable time can pass from nectar uptake to take-off, how patterns affect the take-off decision of flower pollinators could have a distinct impact on the overall handling time of flowers with and without patterns.

To reveal the mechanisms by which flower patterns affect the efficiency of flower handling in insect pollinators, we characterised all stages of flower interaction in a model pollinator, the buffed-tailed bumblebee, *Bombus terrestris*. Using high-speed videography, we analysed the body orientation and flight speed during approach, the position of the touchdown, as well as the walking paths before and after nectar uptake on natural-sized artificial flower models with different pattern types. Our results highlight key mechanisms by which patterns influence flower handling in experienced foragers: first, artificial patterns guided bumblebees to their landing positions, thereby reducing the approach time they required before landing. Second, flower patterns did not reduce the walking time to the nectary, except in a visual conflict configuration. Third, they shaped the inspection paths on the flower after nectar uptake. Together, these effects resulted in a 30% overall reduction in handling time in experienced bumblebee foragers for flowers with radial patterns.

## Results

We quantified the role of flower patterns for different stages of flower interaction: approach flight, landing, and floral exploration before and after nectar uptake in experienced *Bombus terrestris* foragers. Individual foragers were filmed when foraging from flower models of similar size to their grey training models, but with novel colour and pattern features (Fig. 1A). Overall, bumblebees required shorter handling times (from entering an 80 mm radius around the flower to take-off after nectar uptake, excluding the time for nectar uptake) with *cross* patterned flower, compared to *no* pattern and to the *line* pattern (Fig. 1C, generalised linear mixed-effects model (GLMM): p(*no* - *line*)=1.0, p(*no* - *cross*)=0.004, *p(line* - *cross*)=0.033, Table S1). We quantified their head and thorax positions throughout all stages of floral interaction (Fig. 1B,D) upon multiple flower visits, to assess the mechanistic role of flower patterns for the different interaction stages.

### Radial patterns guide approach flight and reduce approach duration

When approaching the flower, the total approach duration (from entering an 80 mm radius around the flower to landing) was significantly shorter for flowers with the *cross* pattern than either the *line* or *no* pattern (Fig. 2A, generalised linear mixed-effects model (GLMM): p(*no* - *line*)=0.98, p(*no* - *cross*)=0.004, *p(line* - *cross*)=0.037, Table S2). As the bumblebees came closer to landing, they reduced their forward velocity (Fig. S2A). This resulted in a significantly lower forward velocity in the 500 ms before touchdown, compared to the 500 ms after take-off when the bees were departing the flower (Fig. 2B, GLMM: *p(approach* - *departure)<0.001*). There were no significant differences in approach velocity during the final 500 ms before touchdown for the different pattern types (Table S3). As for the forward velocity, the rotational velocity of the approaching bumblebees around their yaw axis differed significantly between the approach and departure flights (Fig. 2C, GLMM: *p(approach* - *departure)<0.001*, Table S4). Bumblebees kept a more stable body angle to the flower during approach, while they rotated more freely around their yaw axis upon departure. There were no significant differences in either approach or departure rotational velocity for the different pattern types (Table S3).

**Fig. 2.**
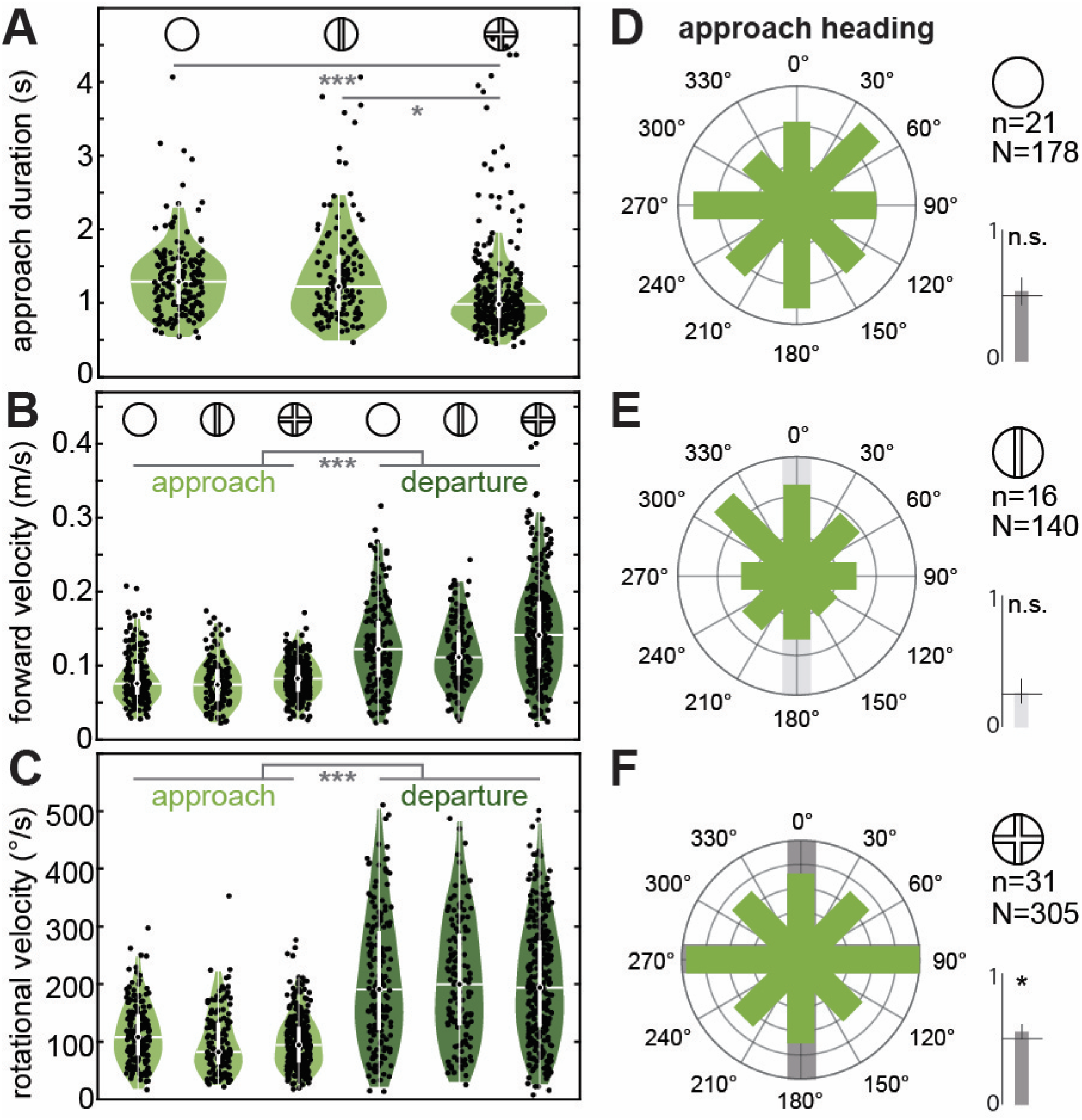
Approach and departure flight characteristics with different flower patterns. **A** shows the total approach duration, calculated as the time from entering a radius of 80 mm from the flower centre to landing, **B** depicts the average forward velocity and **C** the yaw rotation velocity of the *approach* and *departure* flight paths in a 500 ms window before landing and after take-off, respectively, for the three pattern types (*no* pattern, *line* pattern, *cross* pattern). Dots show data from individual flight paths (N as in **D**,**E**,**F**), including multiple trials for individual bumblebees (n as in **D**,**E**,**F**). The asterisks depict the statistical results of a generalised linear mixed-effects model (GLMM, see methods, Table S2–4). Only significant results of the pairwise comparisons of the *pattern-type* and *foraging-time* factors are shown. **D,E,F** Show circular histograms of the average heading angle of each bumblebee’s body axis (defined by the head and thorax position) in the 500 ms approach flights for the three pattern conditions. All data from conditions with patterns was re-orientated so that the patterns aligned with the cardinal axes (the *line* pattern always aligns with the Y-axis), while the *no* pattern condition remained in the original configuration. Thus, in the *no* pattern condition the cardinal axes represent the axes of the video camera’s field-of-view. The heading angles were binned into 8 sectors, of which 2 contain the pattern directions for the *line* pattern (0 and 180°, **E**) and 4 contain the pattern directions for the *cross* pattern (0, 90, 180 and 270°, **E**), as indicated by the grey shades. The statistical results of a generalised linear mixed-effects model implementing a binomial comparison of heading angles (see methods, Table S10) that fall into pattern vs no-pattern sectors are shown as bar graphs next to each histogram. These graphs depict the mean and confidence intervals of heading angles that fell into pattern sectors, relative to the probability of landing in these sectors randomly.

We next assessed whether the rather stable body angles of the bees during approach flight (Fig. 2C) aligned with the flower patterns. We therefore averaged the body angle over the final 500 ms of flower approach and analysed which of the 8 angular sectors of the flower it fell within, where 2 and 4 of the sectors included the pattern in the *line* and *cross* condition, respectively (Fig. 2D-F). With the radial *cross* pattern, the approach heading angles were significantly more frequently in the pattern sectors (Fig. 2F, dark grey shades) than in the background sectors (GLMM with binomial distribution: p=0.026, Table S10). This suggests that the bees used the patterns to control the heading of their approach flight. This was not the case for the approach heading angles in the *no* (Fig. 2D, p=0.277) and *line* pattern conditions (Fig. 2E, p=0.373). Importantly, the approach heading angles in the *no* pattern condition did not reveal any clusters (Fig. 2D), suggesting there were no preferred approach directions, as might have been caused by features of the foraging arena (such as the position of the room’s window, or the entrance to the arena from the hive). Thus, compared to *no* pattern, the duration of the approach flight was significantly reduced with the *cross* pattern, combined with an alignment of the bumblebees’ heading axis with the pattern axes during the approach.

### Flower patterns guided the bumblebees’ landing position

Without a pattern, the bumblebees’ landing positions were spread over the entire angular range of the flower, and over the radial extent (Fig. 3A), though with a higher frequency of landings at the edge than at the centre (Fig. S2A). This general strategy to land at the edge of the flower was also visible in the *line* and *cross* conditions (Fig. S2A). However, in the patterned conditions, there was also an increased proportion of landings close to or on the *line* and *cross* patterns (Fig. 3B, C). We therefore analysed whether bumblebees landed more frequently directly on the patterned area than on the background. Indeed, a significantly higher proportion of bumblebees landed directly on the pattern area for the *cross* pattern (Fig. 3C, GLMM with binomial distribution: p_*cross*_<0.001), with a marginally significant effect for the *line* pattern (p_*line*_=0.096, Table S10). Yet, many bumblebees did not land directly on the patterns, but very close to them. To quantify this effect, we analysed the frequency of landing in the pattern sectors of the flower. For the *line* and *cross* patterns, the landing positions fell significantly more frequent within the pattern sectors than non-pattern sectors (Fig. 3E,F, p*_line_*<0.001, p_*cross*_<0.001, Table S10). Without a pattern, there were no clusters in their landing positions, and there was no increased frequency of landing points in the four cardinal pattern sectors (p*_no_*=0.884). This strongly suggests that the bumblebees used the patterns as landing targets, in particular at the outer rim of the flowers.

**Fig. 3.**
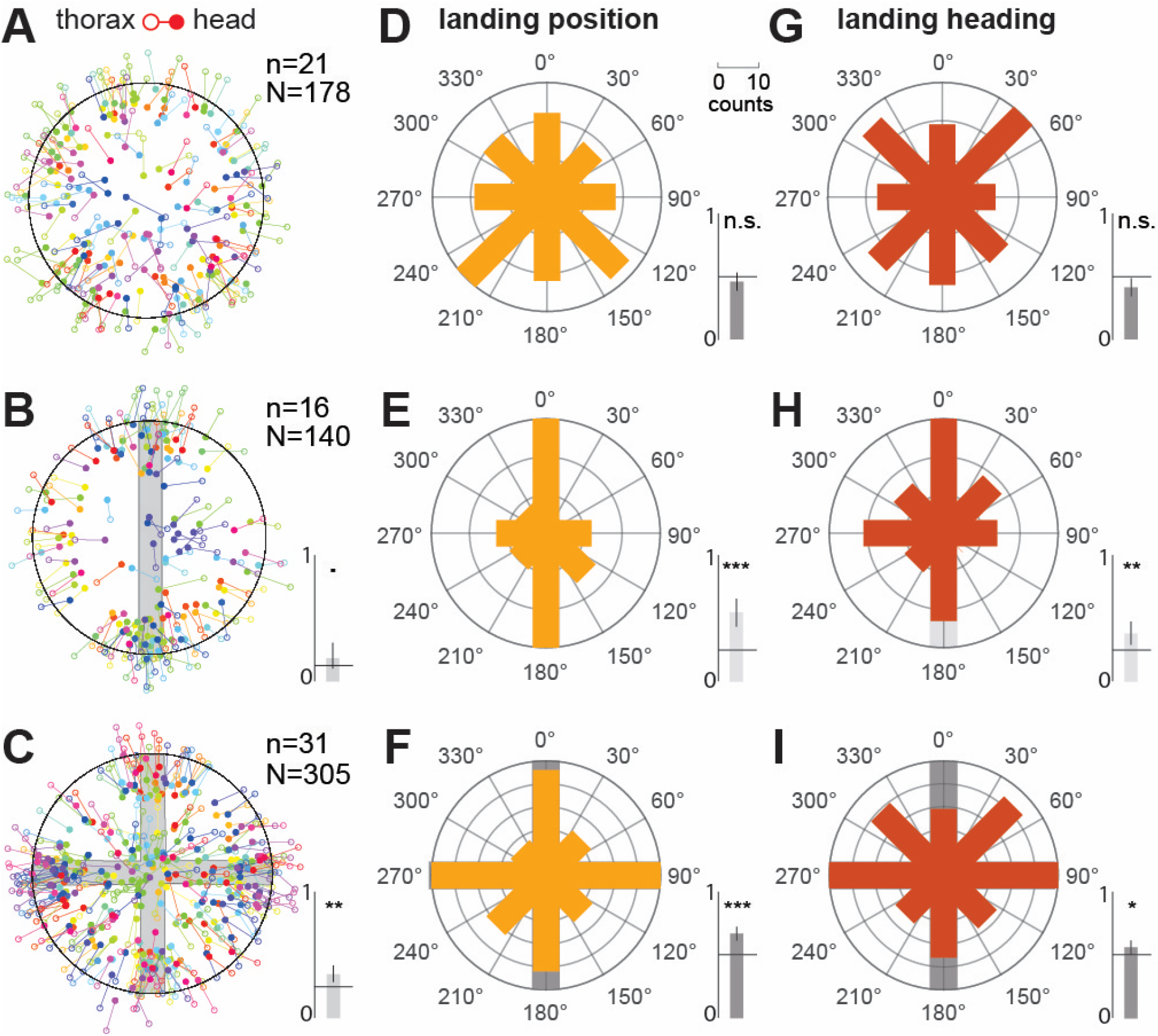
Landing headings and positions with different flower patterns. **A**, **B**, **C** The head (filled circle) and thorax (open circle) position of each bumblebee upon landing is shown for each pattern (indicated in grey). The colours mark individual bumblebees (n), which approached the flowers in multiple trials (N). The inset bar graphs in **B** and **C** depict the results of a statistical test (GLMM with binomial distribution, see Methods, Table S9, 10) to assess whether the bumblebees’ were more likely to land directly on the patterned area than on the background. Depicted are the mean and confidence interval of the probability of a pattern landing, as well as the prediction for random choices (line), which equals the proportion of the flower surface covered by the respective patterns. **D**, **E**, **F** Circular histograms of each bumblebee’s angular landing position (orange) and the orientation of their body axis (defined by the head and thorax position) upon landing (**G**, **H**, **I**, red). The grey shades indicate the pattern sectors in the respective histograms. The inset bars graphs show the statistical results of two GLMMs (see Methods, Table S9, 10) testing the frequency of landing positions and heading angles in the pattern sectors.

We further checked if the bees aligned their body axis with the pattern axis upon landing, so that they landed on or close to the pattern while facing the centre of the flower. Alternatively, bees might have targeted the pattern for landing, but approached it from different angles not aligned with the pattern axes (keeping the pattern perpendicular upon approach, for example), particularly with the *line* pattern. Indeed, for the *line* and *cross* pattern, the frequency of body axis angles aligned with the pattern axes was significantly increased (Fig. 3H,I, p*_line_*=0.001, p_*cross*_=0.015, Table S10). While there were notable exceptions (see Fig. 3B, C), there was a significant population strategy to land on or close to the patterns aligned with the pattern axes, predominately at the edge of the flower.

### The radial extent of flower patterns was crucial for landing guidance

While the bumblebees clearly used the *line* and *cross* patterns to determine their angular landing position and body orientation on the flower, these experiments could not delineate how they chose the distance from the centre at which to land. Since both the *line* and *cross* pattern extended to the rim of the flowers, the bees might either have used the edges of the patterns at the rim as landing targets, or only used their general orientation to align their body axis, choosing their subsequent landing eccentricity independent of the pattern, preferably at the flowers’ edge. To disentangle these effects, we tested a new group of bumblebees with two variations of the *cross* pattern in addition to the original version (Fig. 4A): a *central cross* pattern with 10 mm long arms that extended across only half the flowers’ radius (Fig. 4B), and an *outer cross* pattern (Fig. 4C), that consisted of 10 mm long arms in a cross-orientation, starting at the flowers’ edge, thus leaving the centre free.

**Fig. 4.**
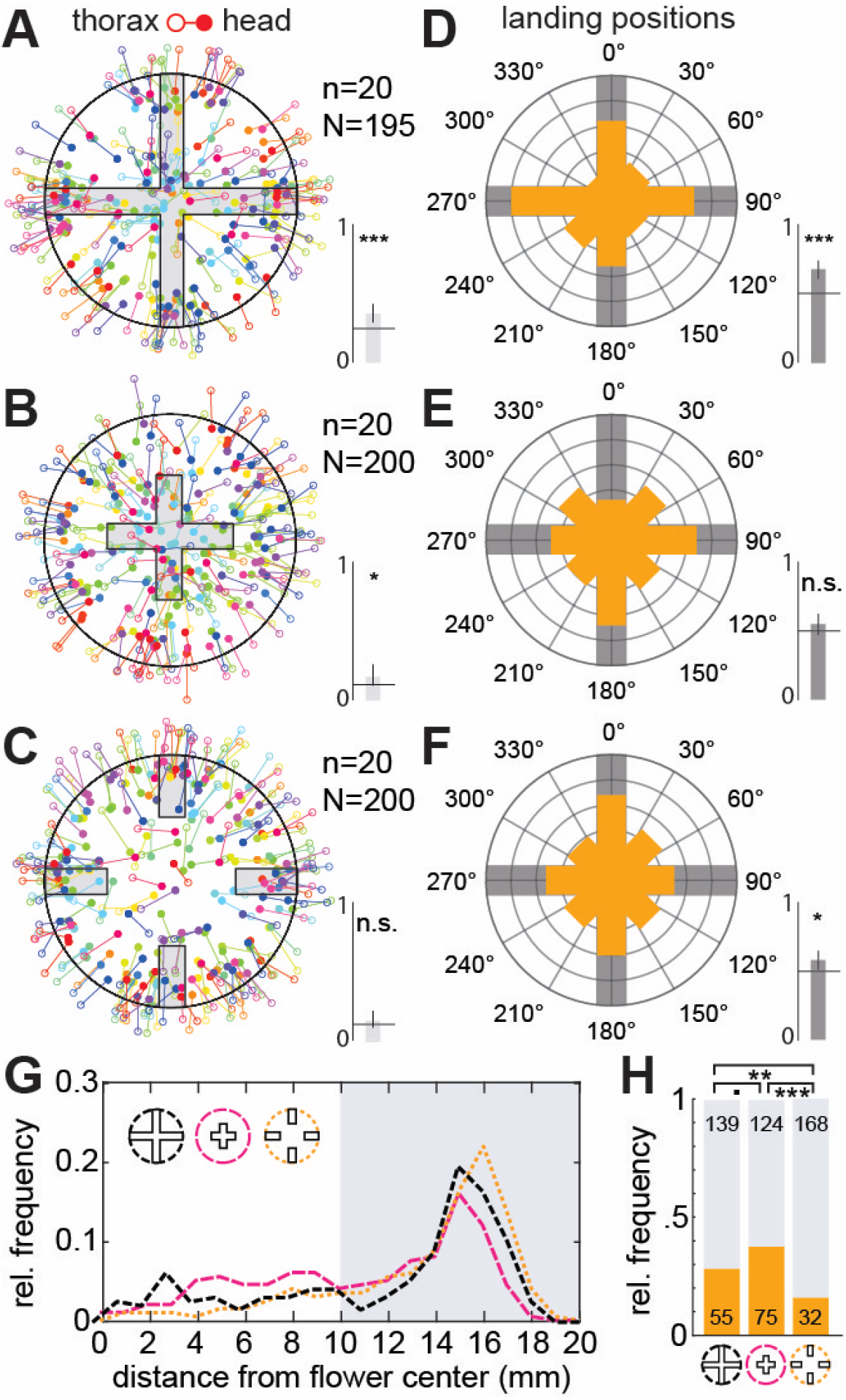
Landing positions with flower patterns of different eccentricity. **A**, **B**, **C** The head (filled circle) and thorax (open circle) position of each bumblebee upon landing is shown for each pattern (indicated in grey). The different colours indicate individual bumblebees (n), which approached the flowers in multiple trials (N). The inset bar graphs in **A**, **B** and **C** depict the results of a statistical test (GLMM with binomial distribution, see Methods, Table S9, 10) to assess whether the bumblebees’ landed directly on the pattern or on the background. Depicted are the mean and confidence interval of the probability of a pattern landing, as well as the prediction for random choices (line), which equals the proportion of the flower surface covered by the respective patterns. **D**, **E**, **F** Circular histograms of each bumblebee’s angular landing position (orange). The grey shades indicate the pattern sectors in the respective histograms. The inset bars graphs show the statistical results of two GLMMs (see Methods, Table S9, 10) testing the frequency of landing positions and heading angles in the pattern sectors. **G** The relative frequency of landings for the three pattern types with respect to the distance from the flower centre, binned at 1 mm intervals. The shaded background highlights two groups for further analysis in **H**: landings closer than 10 mm to the centre and landings further than 10 mm. **H** The number of landings in the inner 10 mm (orange) and outer 10 mm (grey) of the flower was compared for all pattern types (Fisher’s exact test, • = 0.05, * < 0.05, ** < 0.01, *** < 0.001).

With respect to the angular landing position, only the *cross* and *outer cross* pattern resulted in a significantly increased frequency of landings in pattern sectors (Fig. 4D,F, p_*cross*_<0.001, p*_outer cross_*=0.017), while the frequency of landing positions aligned to the inner cross pattern did not differ from chance (Fig. 4E, p*_inner cross_*=0.131). The 4 mm wide arms of the inner cross pattern were still well within a bumblebee’s threshold for spatial resolution: with a spatial resolution of 1° in their frontal visual field (Taylor et al. 2019), a bumblebee could resolve the pattern arm at a distance of 22.9 cm. Thus, we conclude that bumblebees only used patterns that extended to the edges of a flower to guide their angular landing position, but not patterns that were restricted to the flower’s centre, despite their clearly visible directional components.

We next assessed how the position of the patterns’ edges affected the bees’ control of landing eccentricity - the distance from the flower centre. The majority of bumblebees landed at the flowers’ edge (in the outer 10 mm) for all conditions (Fig. 4G). Nevertheless, there was a clear effect of flower pattern on landing eccentricity: the proportion of bees that landed in the centre was significantly higher for the *inner cross* pattern than the *outer cross* (Fisher’s exact test, p<0.001), and marginally significantly higher than for the *cross* pattern (p=0.054). Furthermore, the proportion of bees landing on the outer rim was also significantly higher for the *outer cross* pattern than the *cross* (p=0.003). Thus, despite the prominent contrast of the flower’s edge to the background serving as a strong visual cue for landing control, patterns on the flower did play a role in determining the eccentricity of landing positions.

### Patterns guided movements on the flower

We next investigated whether the flower patterns we presented guided the bumblebees’ approaching walks to the nectary after landing. After landing, the overwhelming majority of individuals walked straight to the presented food reward in the centre of the flower (Fig. 5A). This was visible in short straight approach walking tracks from the landing position to the flower centre (Fig. 5B,C). The duration and tortuosity of the tracks did not differ between the *no*, *line* and *cross* pattern conditions (duration: Fig. 5B, GLMM: p_no-*line*_=0.719, p_no-*cross*_=0.542, p_line-*cross*_=0.983, Table S6, tortuosity: Fig. 5C, p_no-l*ine*_=0.986, p_no-*cross*_=0.854, p_line-*cross*_=0.776, Table S5). There was also no significant difference in approach duration between the *cross* and *inner cross* patterns (Fig. S4A, Table S7). Thus, the bumblebees made their way to the nectar reward on the straightest path possible, irrespective the presence or type of the flower patterns. The walking paths of bumblebees with the *outer cross* pattern were the only exception to this trend,, requiring significantly longer to reach the nectary than for all other cross patterns (Fig. S4A, p_cross-o*cross*_=0.036, p_icross-o*cross*_=0.031, Table S7). The tortuosities of these paths were similar to the other cross pattern conditions (Table S8). This indicates that the bees did indeed rely on some of the visual patterns to guide their approach walk to the nectary.

**Fig. 5.**
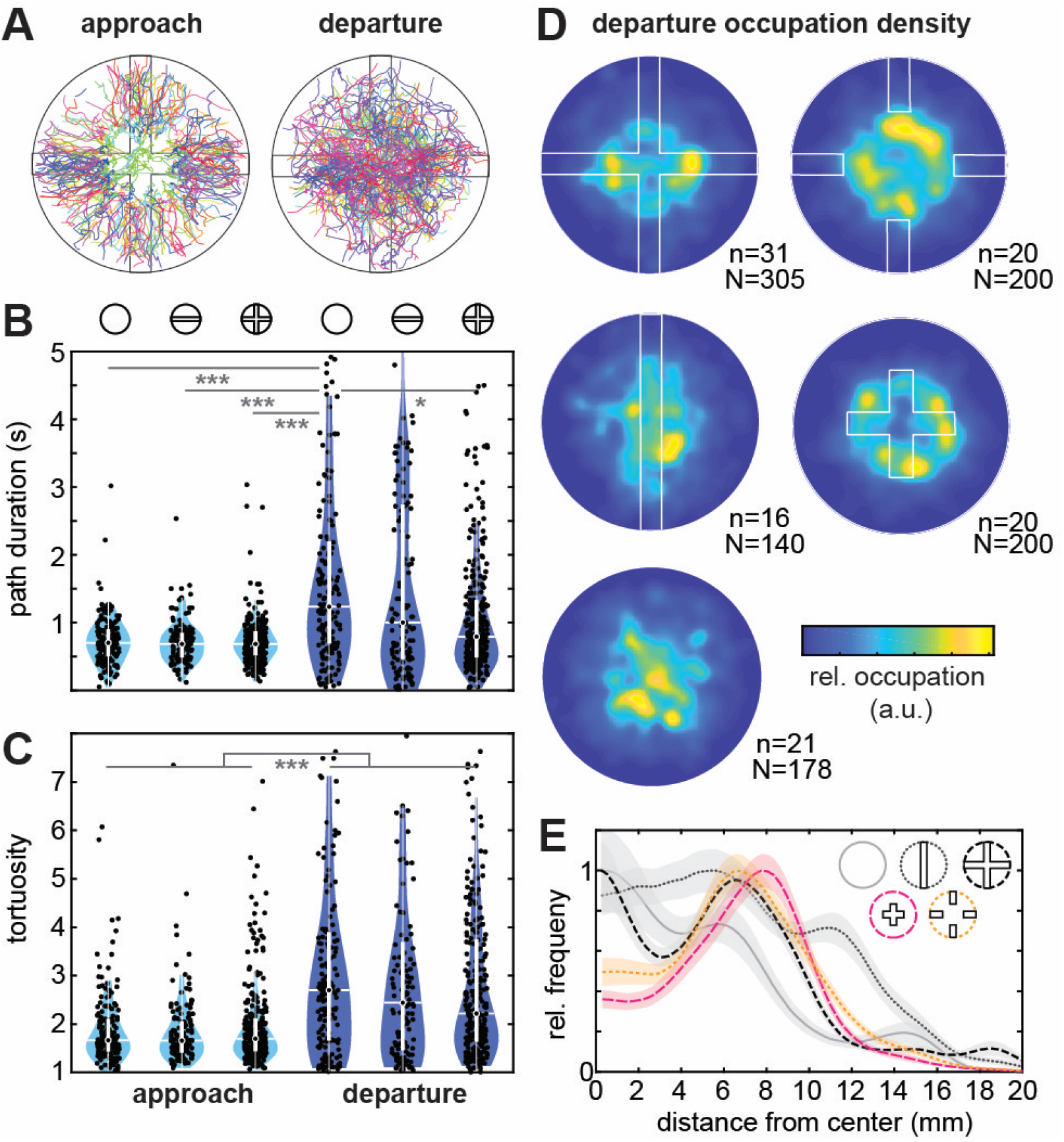
Walking paths on the flower with different patterns. **A** The walking paths of individual bumblebees after landing until first proboscis contact with sucrose (*approach*) and after drinking until take-off (*departure*). The different colours indicate individual bumblebees (n) with multiple trials (N, as indicated in **D**). **B,C** Path duration (**B**), and tortuosity (**C**) of the *approach* and *departure* walks, respectively, in three pattern conditions. Dots show data from individual flight paths (N), including multiple trials for individual bumblebees (n, see **D**). The asterisks depict the statistical results of a generalised linear mixed-effects model (GLMM, see methods, Table S5, 6). Only significant results of the pairwise comparisons of all conditions (pattern type, flight type) are shown. **D** Heat maps show the average relative occupation density of the bumblebees’ departure walks on the different pattern types. These were generated from the normalised occupation density generated from each individual bumblebee, where each position per timeframe contributed one entry to the individual’s occupational density, which was then averaged across all animals. **E** The relative frequency of occupation of departure walks along different radial distances from the flower centre. These frequencies were generated as a radial average of the individual occupational densities, and subsequently averaged. The graphs show the mean and standard deviation for the different patterns.

The movement tracks of the animals after nectar intake were much more variable than during approach, ranging from a few steps before take-off to long explorative paths on the flower (Fig. 5A). This resulted in a broader range in duration and tortuosity of the departure walks than approach walks (Fig. 5B,C). Indeed, approach and departure were a significant factor in walk tortuosity (GLMM: p_approach–departure_<0.001, Table S5), and in walking path duration (GLMM: p_approach–departure_<0.001, Table S6). While the tortuosity differed significantly between the approach and departure portions of all the paths in all pattern conditions (Table S5), this was only the case for some of the patterns, approach/departure and flower colour interactions, in particular those with *no* pattern (Table S5), where the departure walks of the *no* pattern were significantly longer.

Given that flower patterns influenced not just the duration, but also the straightness of the departure walks, we next investigated whether the patterns also influenced where the bees walked on the flower. Indeed, the areas the bumblebees frequented most during their departure walks differed across pattern types (Fig. 5D): with *no* pattern present, the highest occupation density was in the centre of the flower, while with the *line* pattern, the bees occupied a more elongated area along the pattern. With all three *cross* pattern types, the occupied area formed a ring shape around the centre. To compare the spatial structure of the occupation density further across pattern types and individual animals, we calculated the relative radial occupation density per individual, and averaged it across individuals (Fig. E). This analysis shows the most frequented positions on the flower with respect to the eccentricity from the centre. The ring structure visible with the *cross* patterns in the 2D occupation maps (Fig. 5D) is represented in this analysis as a distinct peak at approximately 8 mm distance from the centre – and reveals that the bumblebees frequented the edges of the *central* and *outer cross* more frequently than other regions of the flower. They also frequently explored this area with the *cross* patterns in addition to the centre of the flower, even though the cross arms extended to the rim of the flower. In contrast to the *cross* patterns, the highest occupation density with *no* pattern was in the centre of the flower. Taken together, the analysis reveals that unlike for approach walks, which were directed straight to the centre of the flower, the patterns played a distinct role in shaping the duration and spatial structure of departure walks.

## Discussion

In this study, we demonstrated that flower patterns increase the foraging efficiency of experienced insect pollinators by decreasing the flower handling time at several stages of their flower visit. By analysing their flight and walking behaviour for the entire flower interaction, we could highlight the mechanisms underlying this improved efficiency. Contrary to previous reports from naïve foragers, we found no significant impact of radial patterns on the walking paths of bumblebee foragers to the nectary. The strong reduction in handling time resulted from the patterns’ role in guiding the bumblebees’ approach flight and landing, as well as their walking paths on the flower before departure.

### The role of patterns for approach and landing guidance

Flower patterns are thought to have evolved to improve foraging efficiency by reducing handling times of the pollinator during a flower visit (Dinkel & Lunau 2001, Goodale et al. 2014, Goyret 2010, Goyret & Kelber 2012, Johnson & Dafni 1998, Lawson et al. 2017, Leonard & Papaj 2011, Waser & Price 1985) – in addition to their role in flower selection (de Ibarra et al. 2015, Orbán & Plowright 2014, Stöckl & Kelber 2019). However, the major role of flower patterns in flower handling is commonly assumed to indicate the position of the nectary, and specifically to guide direct approaches to the nectary after landing on a flower. Our results highlight a crucial and generally overlooked function of flower patterns: efficient guiding of the approach flight and landing on the flower, irrespective of the position of the nectary. The 31% reduction in approach duration facilitated by radial patterns (Fig. 2A) shows that even though the bees were not guided directly to the nectary, as they landed mainly on the edge of the flower (Fig. S3), this effect provided a substantial improvement of foraging efficiency.

We could further pinpoint how the patterns contributed to a reduction in approach flight duration: they served as guides for both the angular bearing (Fig. 3) and the eccentricity of the landing position (Fig. 4). Thus, by providing uniquely identifiable landing positions, they may reduce the inspection time approaching bumblebees require to initiate landing, consequently reducing their overall approach flight time. The observed pattern-based control of eccentricity occurred despite a strong preference of the bees to land at the edge of the flowers in all pattern conditions, likely aided by the strong visual contrast between the flower and the grey background. This confirms previous observations on much larger artificial flowers (Free 1970, Manning 1956), and provides strong evidence that these guidance effects also occur on biologically relevant flower sizes.

The effects of patterns on angular landing positions have not been observed previously. They were closely related with the alignment of the bees’ body axis to the flower patterns during approach flight (Fig. 2C-E) and landing (Fig. 3G-I). This suggests that the bumblebees used the pattern arms as cues to stabilise their approach flight before touchdown. As has been shown in honeybees (Rusch et al. 2021) and flies (Maimon et al. 2008), many insects fixate elongated targets, for example stripes, in their central visual field when they move. The arms of the flower patterns we presented could have served as such fixation targets.

However, we observed distinct differences in the bumblebees’ responses to the different pattern types: while the *line* pattern resulted in pattern-related angular landing positions (Fig. 3E) and body orientations during landing (Fig. 3F), it did not induce an alignment of the body orientation with the pattern, or a significant reduction in the approach duration compared to the *cross* pattern (Fig. 2A). One major difference between the *line* and *cross* pattern is that the *cross* pattern provides translational optic flow and alignment cues at the same time: if an approaching bumblebee aligned their body axis with one arm of the *cross* pattern, the respective cross-bar of the pattern generated translational optic flow. The *line* pattern, on the other hand, either provided translational optic flow cues (if bees approached the pattern perpendicularly), or alignment cues (if they approached in parallel). Previous studies on honeybees demonstrated the importance of translational optic flow cues to control the flight speed upon landings on horizontal surfaces (Srinivasan et al. 2000) and of motion cues to identify edges suitable for landing on horizontal targets (Lehrer & Srinivasan 1993). Thus, the bees benefitted from aligning with the arms of the *cross* pattern and landing at their contact with the flowers’ edge in all aspects of their approach flight and landing. In contrast, an alignment with the *line* pattern would have only been partly beneficial, as optic flow cues for flight control would have been generated most strongly by a perpendicular approach. In addition, the radial pattern generated stronger expansion cues upon approach than the line pattern – which are important for landing control in bees (Baird et al. 2013). These differences in approach and landing guidance between pattern types strongly suggest that the bumblebees used patterns both for flight stabilisation, in addition to selecting a unique landing position.

### The role of flower patterns during landing

Many previous studies have suggested that flower patterns serve as ‘nectar guides’ (Dinkel & Lunau 2001, Goodale et al. 2014, Goyret 2010, Goyret & Kelber 2012, Johnson & Dafni 1998, Lawson et al. 2017, Leonard & Papaj 2011, Sprengel 1793, Waser & Price 1985), thus guiding pollinators to the nectary (or to the pollen, in the case of ‘pollen guides’) by indicating their respective positions. Our results do not support this notion for experienced bumblebee foragers. One might have expected that the bumblebee foragers would use the *cross* patterns in particular to land at the centre of the flower, where the sugar reward was presented in all of the experiments and during training. The foragers participating in the experiments had experienced the nectary in the flower centre at least 30 times on the grey patternless training flowers before being confronted with the test patterns, and could also see the sugar reward placed openly as a small drop on the flower surface (see Methods). Nevertheless, the majority of foragers landed at the edge of the flower (Fig. 3, 4 & S3), and did not use the flower patterns as indicators to land directly at the nectary (Figs. 4G & S3A).

These findings are supported by previous observations, which showed that honeybees and bumblebees did not land on the convergence point of several lines presented on artificial flowers. Instead, bees landed all along the lines in irregular orientations (Free 1970, Manning 1956), suggesting that they did not interpret the spatial layout as nectar guides, but were attracted by their colour or contrast of the pattern in general. This is in line with our results, where the bumblebees landed significantly more often on the edge of flowers with outer crosses, rather than at their convergence point in the centre (Fig. 4), even though the bees had already learnt to find the nectary in the flowers’ centre during training. Thus, while guiding the foragers’ approaches and landings on the flower in general, bumblebees did not use flower patterns to approach the nectary in the centre of the flower directly.

### The role of flower patterns as guides to the nectary – walking approach

Our results did not provide evidence that flower patterns improved the efficiency of nectary discovery once the bumblebees landed, as there was no difference in walking time across the *no*, *line* and *cross* pattern conditions (Fig. 4B). Instead, for all conditions except the *outer cross*, the bumblebees walked in rather straight paths from their landing position to the food reward in the centre of the flower (Fig. 4A). At first sight, this contradicts a previous study with comparably sized and patterned artificial flowers, which described a significantly reduced time until the discovery of the nectary after landing when patterns were present, suggesting a guidance function of the patterns (Leonard & Papaj 2011). However, this discrepancy might be due to a different role of the flower patterns in the respective foraging setting: (Leonard & Papaj 2011) presented the patterns in the context of a learning task, where only a subset of flowers with patterns, or respectively without, was rewarded. Thus, differences in search times might be due to the time required for flower identification, rather than nectary location – a well-demonstrated function of flower patterns during search behaviour by flower visitors (Dafni et al. 1997, Hansen et al. 2011, Heuschen et al. 2005, Johnson & Dafni 1998, Kelber 2002, Lunau et al. 2009). In our task, all flowers were rewarded, so that the flower patterns did not indicate a reward, but might act as potential landing and nectar guides.

Moreover, our experiments also differed from those of (Leonard & Papaj 2011) in the bumblebees’ experience of the flower’s configuration, and how they learned to locate the nectary. (Leonard & Papaj 2011) showed that foraging experience drastically reduces walking time to the nectary in both flower types, with and without patterns. They furthermore showed that the influence of patterns on the approach time depended on which patterns had been rewarded during training. Bees that were initially rewarded on un-patterned flowers did not decrease their approach time with patterned flowers, while the opposite was the case for the reversed order of presentation. Without patterns during learning, bees might thus learn the nectary configuration relative to the flowers’ shape, but if patterns are present, bees might learn the nectary position relative to the patterns, and thus require longer to locate the nectary without them. In our study, the foragers were extensively trained to pattern-less grey flowers, and likely learnt the nectary configuration. This would explain that the approach walking times did not differ between the patterned and un-patterned flowers (Fig. 4), as they might have mainly relied on their knowledge about the flower configuration. However, the bumblebees in our experiments did not entirely ignore the patterns: for the *outer cross* pattern, which created high-contrast visual cues separate from the nectary position, approach walking times were significantly longer than for all other patterns (Fig. S4). This suggests that experienced bees relied on their experience of the pattern configuration even when flower patterns were present, but when patterns provided conflicting cues about the position of the nectary, bees were distracted during their nectary approach.

Thus, we conclude that the role of flower patterns as walking guides to the nectary strongly depends on the previous foraging experience of bumblebees, and on how the pattern cues combine guidance information with the learned flower and nectary configuration.

### Flower patterns influence departure decisions

While bumblebees did not reach the nectary faster on flowers with or without patterns, we found that patterns significantly influenced departure decisions: radial patterns reduced the bumblebees’ exploration time on the flower after emptying the sugar reward (Fig. 5A, Fig. S4A). These observations are supported by a previous study (Leonard & Papaj 2011). Our analysis of the bees’ exploration tracks, however, provides an explanation for this finding: the bees’ paths on the flowers were specifically clustered around the contrast edges of the patterns (Fig. 5D,E), suggesting that the flower patterns guided the post-drinking exploration of foragers. A lack of patterns might suggest hidden nectaries or pollen, and thus might encourage an extended search.

Our findings suggests that in future studies, more emphasis should be placed on the post-nectary exploration to better understand the role of flower patterns for foraging efficiency, as it took a considerably larger share of the foragers’ time on the flower than the nectary approach (Fig. 5B, also in (Leonard & Papaj 2011)). Longer exploration times increase the risk of exposure to predators on the flower and reduce the time available to exploit other food sources. Future work may focus on identifying the observed post-drinking exploration patterns in real flowers, which would provide a better understanding of the diverse functions of floral patterns in plant-pollinator interactions.

### What do these experiments suggest about foraging in the natural world?

Based on our laboratory experiments, a number of key predictions can be made about the relevance of flower patterns for insect pollinators at real flowers under natural conditions. Together with insights from comparable studies (Free 1970, Leonard & Papaj 2011, Manning 1956), our results suggest that the role of floral markings for guiding a flower visitor to the nectary after landing strongly depends on its foraging experience. The impact of floral markings drastically decreases with experience, to the point where approach times to flowers with and without patterns may not differ. Honeybee and bumblebee foragers perform several hundreds to a thousand flower visits during a day (Abrol 2012, Pasquaretta et al. 2019) – compared to the dozens of visits on which most laboratory observations are based – and are known to improve their natural flower handling with experience (Laverty 1980). We thus can expect that flower patterns play only a minor role for finding the nectaries while walking on the flower for the majority of their foraging life. Moreover, on a natural flower, sensory cues beyond visual ones may guide the bee to the nectary after landing: the shape of the flower, as well as mechanosensory, olfactory and humidity cues, thus further diminishing the potential importance of flower patterns for nectary guidance after landing. Interestingly though, when flower patterns provided conflicting information about nectary location with regard to the learned configuration, they resulted in increased exploration times even in experienced foragers (Fig. S4A). Such scenarios might be employed by some flowers to trick insects into suitable positions for pollination (Dafni 1984), or deceit pollinators about the availability of nectar (Thakar et al. 2003). Mismatched visual cues and nectary location may also occur in ornamental flowers, where the floral markings have been altered from their natural configurations.

Importantly, our results demonstrate that even for experienced foragers who have learned the position of the nectary on the flower, flower patterns are important to guide their approach flight and landing on the flower, thus significantly reducing handling time. Since approach guidance and landing control cannot be supported by other sensory modalities (except for mechanosensation in the legs and antennae upon contact with the flower (Reber et al. 2016)), we expect this effect also to be effective in a natural setting. Indeed, we expect approach guidance and landing control to be of even higher importance than in the laboratory, as flowers in nature are encountered at different heights, tilt angles, and vary in size and shape (Dafni & Kevan 1997, Dafni et al. 1997), making it harder to transfer learned approach-and-landing strategies between flowers, and therefore relying more on sensory inputs. Additional challenges during flower approach and landing are obstacles such as foliage in the potential approach paths, changing lighting conditions, as well movement of the pollinators and flowers induced by wind (Chang et al. 2016).

While the exclusively horizontal presentation of our artificial flowers was not representative for all flower types visited by bumblebees, their size and general shape fell well within the natural range of flowers bumblebees typically forage from (Dafni & Kevan 1997, Dafni et al. 1997). Moreover, choice experiments suggest that bumblebees prefer horizontally arranged flowers (Makino 2008), from which they also learn cues more flexibly (Wolf et al. 2015), so that this type of flower is highly relevant for bumblebees. The colour contrast and spatial resolution of the patterns in our experiments were designed to ensure reliable detection by the bumblebees. It is currently unclear whether this is comparable to most natural flower patterns of bumblebee pollinated flowers, as the spatial colour patterns of most insect pollinators’ host flowers remain largely unquantified (Garcia & Dyer 2021), (but see (De Ibarra & Vorobyev 2009, Vorobyev et al. 1997)). With the advent of new multispectral imaging techniques (Garcia et al. 2014, Lunau et al. 2021, Tedore & Nilsson 2019), in the coming years a higher-throughput quantification of flower patterns will likely provide a better understanding for how natural flower patterns compare to the stimuli used for laboratory experiments, enabling more natural stimulus designs for presentation under controlled laboratory conditions.

### Conclusion

Our results reveal previously overlooked guidance functions of flower patterns that contribute significantly to foraging efficiency. We demonstrate how patterns guide the approach flight and landing control of bumblebees on flowers, and which pattern features play a role in this guidance. Moreover, we highlight the role of patterns in the decision to depart or to continue flower inspection after nectar uptake. These insights provide a novel focus for the study of flower-pollinator interactions in the lab and in their natural environment and open up new questions on how deceptive patterns or altered displays on ornamental flowers impact foraging efficiency in insect pollinators.

## Materials and Methods

### Animals

Colonies of *Bombus terrestris* were obtained from commercial breeders (Koppert, Berkel en Rodenrijs, Netherlands and Biobest, Westerlo, Belgium). The colonies were housed in a two-chamber wooden nesting box (28 cm×16 cm×11 cm), which provided one chamber for nest-building and one for feeding. Before experiments, the bees were fed ad libitum with APIinvert (Südzucker, Germany), a 70% sucrose solution, and with pollen grains (Bio-Blütenpollen, Naturwaren-Niederrhein, Germany).

### Setup and Experimental Procedure

For free flight experiments, the nesting boxes were connected to a flight arena (120 cm × 100 cm × 35 cm) via an acrylic tube. The arena was constructed from plywood, with a UV-transparent acrylic lid. It was illuminated from directly above by 2 fluorescent tubes (Osram, Biolux, L58W/965) connected to an electric ballast, as well as indirect light through the room’s windows, resulting in a light intensity of 510 lx at the bottom of the arena pointing upwards, with the spectrum shown in Fig. S1A. The floor of the arena was covered by grey cardboard (Mi-Taintes #122, von Canson SAS, Annonay Cedex, France), and the same cardboard was used for the training feeders (see below). Bumblebees visiting the flowers were recorded using an ELP USB camera (2.0 Megapixel (1080p), Ailipu Technology), operated at 60 fps with 1280 × 720 pixel using the software ContaCam 7.9.0 beta7 (Contaware) in motion detection mode for automatic video acquisition. The camera was directly attached to the acrylic lid to minimise reflections from the overhead lighting, and positioned above the respective stimuli that were imaged.

### Stimuli

All artificial flowers were constructed from round paper or cardboard cutouts of 40 mm diameter, mounted on 10 mm high dark grey platforms of 40 mm diameter. The grey training stimuli were constructed from the same cardboard used for the floor of the arena, while the coloured test stimuli were printed on un-bleached paper (“Classic White”, Steinbeis, Glückstadt, Germany) using a laser colour printer (C3325i, Canon, Germany). All stimuli were subsequently laminated with a matt foil (S-PP525-22 matt, Peach, PRT GmbH, Tannheim, Germany). The yellow colours were printed as C=0%, M=0%, Y=100%, K=0%, and the blue as C=70%, M=15%, Y=0%, K=0%. The relative reflectance spectra of the resulting colours are shown in Fig. S1B. We used three different main categories of stimuli: blue and yellow artificial flowers without patterns (*no* pattern), and artificial flowers with a *line*, and a *cross* pattern, which were either presented in yellow on blue background, or vice versa. The arms of all lines or crosses were 4 mm wide, and the central point of the cross positioned at the flowers’ centre.

In this study, we performed two experiments, in which different pattern types were shown to separate groups of bumblebees. In the first experiment, we presented yellow and blue artificial flowers without a pattern (*no* pattern), a *line* and a *cross* that extended to the rim of the flower. The second experiment consisted only of cross-type stimuli: the same *cross* as in the first experiment, an *inner cross* with arms of 20 mm length from the centre of the flower, and an *outer cross*, which consisted only of 20 mm-long arms extending from the rim of the flower to the centre, but lacking the central part of the cross (Fig. S1C). In each experiment, we presented the two possible colour combinations of each of the three stimuli to each bumblebee, resulting in six different pattern conditions per individual.

### Experimental Procedure

*B. terrestris* foragers were selected for experiments by tracking their foraging activity in the flight arena on grey training flowers covered generously with 30% sugar solution. They were marked on the thorax with individual number tags. In each experimental session, before being presented with the test stimuli, five grey training flowers were positioned in the arena in a random arrangement, with one placed directly under the central camera. They were prepared with 10 μl of 50 % sugar solution in the centre of the flower. The bees were allowed three foraging bouts on the grey flowers. Foraging bouts were separated by the forager returning to the colony and comprised between 10 and 20 flower visits each. Each flower was refilled immediately after the bumblebee departed. After the three initial foraging bouts on grey flowers, the first test stimuli were presented. Five flowers of the same type were randomly arranged in the arena, the central one being placed under the camera as for the grey ones, and the same system of sugar reward. Five visits to the central flower were recorded, which were contained within a maximum of three foraging bouts. After that, one foraging bout on grey flowers was conducted, followed by the next pattern type, until all six pattern types were tested. Pattern types were presented in a randomised order. Typically, all data from one forager was collected within a single day. In cases were the forager aborted their foraging activity before all six pattern types could be tested, missing data were collected on the following day.

### Video tracking

We used DeepLabCut (Mathis et al. 2018) to automatically identify the head and thorax position (Fig. 1B; using 4100 frames, merged from 5 different recordings as a training dataset). All videos were manually curated afterwards using the DLTdv7 software (Hedrick 2008) in Matlab 2017b (The Mathworks, Natick, US), to ensure a high precision of tracking throughout the entire video. The flower position (marked by four points on the flowers’ circumference parallel to the cardinal axes) and pattern position (marked on all pattern edges) were indicated in each video using DLTdv as well. Furthermore, the landing position was marked manually, with landing defined as the touchdown of at least the two front legs with no immediate subsequent take-off. We also marked the initiation and end of the drinking phase (indicated by the extension of the proboscis into the sugar solution, and its retraction, respectively), to separate flight and walking trajectories before and after nectar uptake (*approach* and *departure*, Fig. 1C). Data were then further processed using custom-written Matlab scripts.

### Data analysis

From the bumblebees’ approach flight trajectory, we calculated the time from entering an 80 mm perimeter around the flowers’ centre to the bumblebees’ landing (Fig. 2A). We further analysed the approach and departure trajectories 500 ms before landing and after take-off, respectively, to calculate the average forward velocity as the mean projected ground speed between consecutive tracking points (Fig. 2B, Fig.S2A.B), and the rotational velocity as the change in orientation angle of the longitudinal body axis (Fig. 2C, Fig. S2C.D). We determined the longitudinal body axis using the head and thorax tracking points. We furthermore used the 500 ms approach and departure trajectories to calculate the bees’ mean heading direction as the circular mean of frame-by-frame body axis orientations (Fig. 2D-F, Fig. S2E-G). Where patterns were presented, the heading directions were calculated relative to the pattern axes (so that the patterns aligned with the cardinal axes, where the *line* pattern always aligned with the Y-axis, Fig. 2E,F). In the *no* pattern condition, heading directions were analysed relative to the flower’s absolute position in the arena, so that the cardinal axes represent the axes of the video camera’s field-of-view (Fig. 2D). This condition thus provided a control for potential biases in the absolute approach directions.

We sorted the approach and departure headings into 8 evenly spaced sectors (bin centers of 0°, ±45°, ±90°, ±135°, 180°), relative to the pattern’s orientation (or camera axes for the *no* pattern condition): with the *line* pattern, two of the sectors contained the pattern directions (0 and 180°, Fig. 2E, grey shade in histogram) and four with the *cross* pattern (0, 90, 180 and 270°, Fig. 2E, grey shade in histogram). We used the same histogram bins for the landing positions (Figs. 3D-F, 4D-F), as well as for the landing headings (the orientation of the body axis of the bumblebee upon touchdown, Fig. 3G-I). We furthermore quantified the distance of the landing positions from the flowers’ centre (Fig. 4G, Fig. S3A), and calculated the relative frequency of landings inside the perimeter of half the flowers’ radius and outside of it (Fig. 4H, S3B).

Using the touchdown and take-off timestamps, as well as the timestamps of nectar retrieval on the flower, we separated the bumblebees’ walking paths on the flower into approach and departure tracks (Fig. 5A). We calculated the paths’ total duration (Fig. 5B) and the tortuosity (as the paths’ length divided by the distance between its start and end point, Fig. 5C). We furthermore calculated the relative frequency of occupation for each pixel position on the flower, based on the departure tracks of each bumblebee (Fig. 5D). To compare the density at different distances from the flower centre, we summed the occupation density of each bumblebee flower visit over the radial axes of the flower in 1° steps and normalised it to the maximum before averaging across bees.

For visual presentation, we pooled the data obtained from blue patterns on yellow background and yellow patterns on blue background, since the statistical analysis did not reveal any significant interaction of pattern colour with the analysed measures (see *statistical analysis*).

### Statistical analysis

To statistically analyse the effect of pattern type, pattern colour and individual identity on the flight, landing and walking parameters of the bumblebees, we used generalised linear mixed-effects models with R v4.1.2 (R Foundation for Statistical Computing).

To analyse the heading, landing position and landing orientation data, we used generalised linear mixed-effects model with the formula

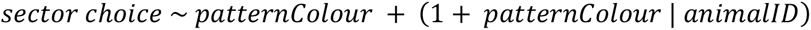

These estimated the fixed-effect of pattern colour on the relative probability of a landing in a pattern or non-pattern sector – scored for each individual flight as 0 or 1, accounting for individual biases and differences in response to pattern colour. For the *no* pattern condition, the sectors containing the cross pattern were identified as part of the target region, to test for any general preference for landings along the cardinal axes of the camera’s field-of-view (and thus also of the flight cage). Thus, the random choice probability for the *no* and *cross* pattern condition was 0.5, and for the *line* condition it was 0.25 (see Table S10).

Since the *patternColour* factor did not significantly reduce model deviance with any of tested parameters (assessed by comparing the deviance of the full and reduced model below using a likelihood-ratio test, Table S9), we combined the data for all colour conditions, and fitted the following model, accounting only for individual biases, for all subsequent analysis:

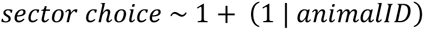

The resulting model (mean and confidence intervals) is shown in each figure as an inset bar graph. The statistical results are summarised in Tables S10.

To analyse the total duration of the approach flight, the forward and rotational velocity of the flight tracks, and the total duration of the walking paths and their tortuosity, we fitted a generalised linear mixed-effects model with the formula

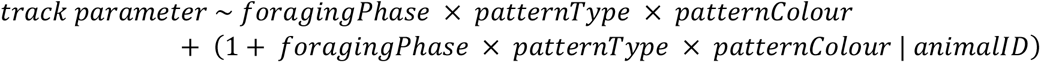

For the analysis of approach duration, the *foragingPhase* (approach, departure) factor was not included in the model, since this analysis was only conducted on the approach data. All data was log-transformed to ensure normality of the residuals.

Where the patternColour factor did not significantly reduce model deviance (see Table S1–8), we combined the data for all colour conditions for the subsequent analysis, and fitted the following model:

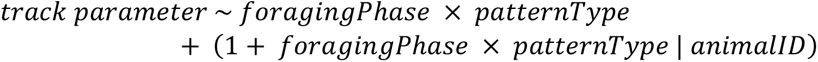

Where the *patternColour* factor significantly reduced model deviance, we always analysed the full model with the factor (including cases where the data is pooled across colour conditions in the figures for better visibility). The statistical results are summarised in Tables S1–8.

## Acknowledgments

We acknowledge funding to A.S. from the Volkswagen Foundation (95490), the PostDoc Plus Program of the Graduate School of Life Sciences at the University of Würzburg, and the Bavarian Academy of Sciences and Humanities.

## Author Contributions

Conceptualization: A.S., J.S.; Methodology: A.S., J.S.; Validation: R.R., A.D., A.S., J.S., J.F.; Formal analysis: R.R., A.D., J.F., A.S.; Investigation: R.R., A.D.; Resources: A.S., J.S.; Data curation: A.S.; Writing - original draft: A.S.; Writing - review & editing: R.R., A.D., J.F., J.S., A.S.; Visualization: A.S., J.F.; Supervision: J.S., A.S.; Funding acquisition: J.S., A.S.

## Supplementary Figures

**Fig. S1.**
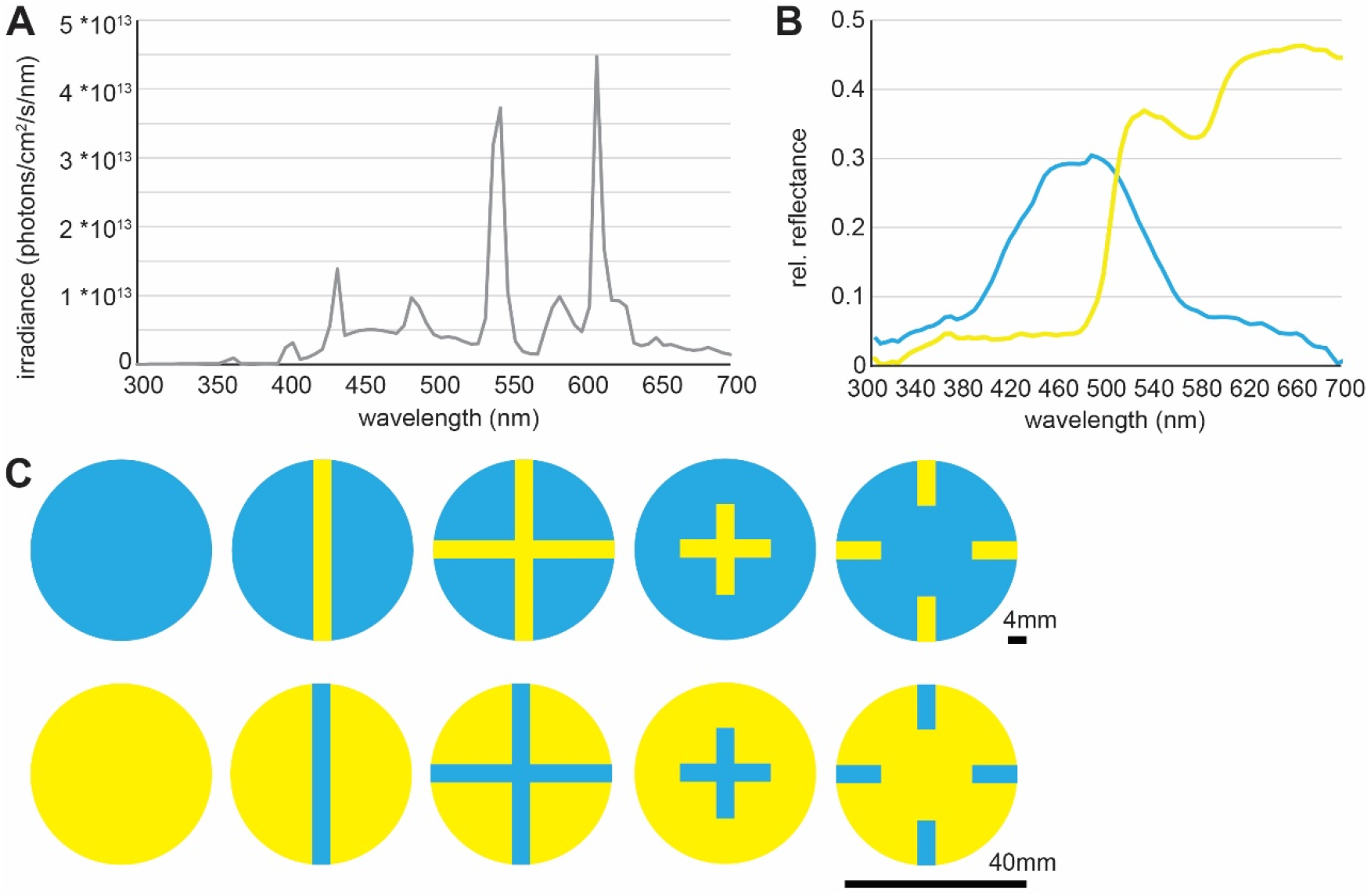
Flower patterns used in this study and their spectral reflectance. **A** Irradiance spectrum of the light sources in the flight arena measured from the floor of the arena pointing upwards. **B** The relative reflectance spectra of the colours of the test stimuli. **C** The different types of flower pattern used in this study.

**Fig. S2.**
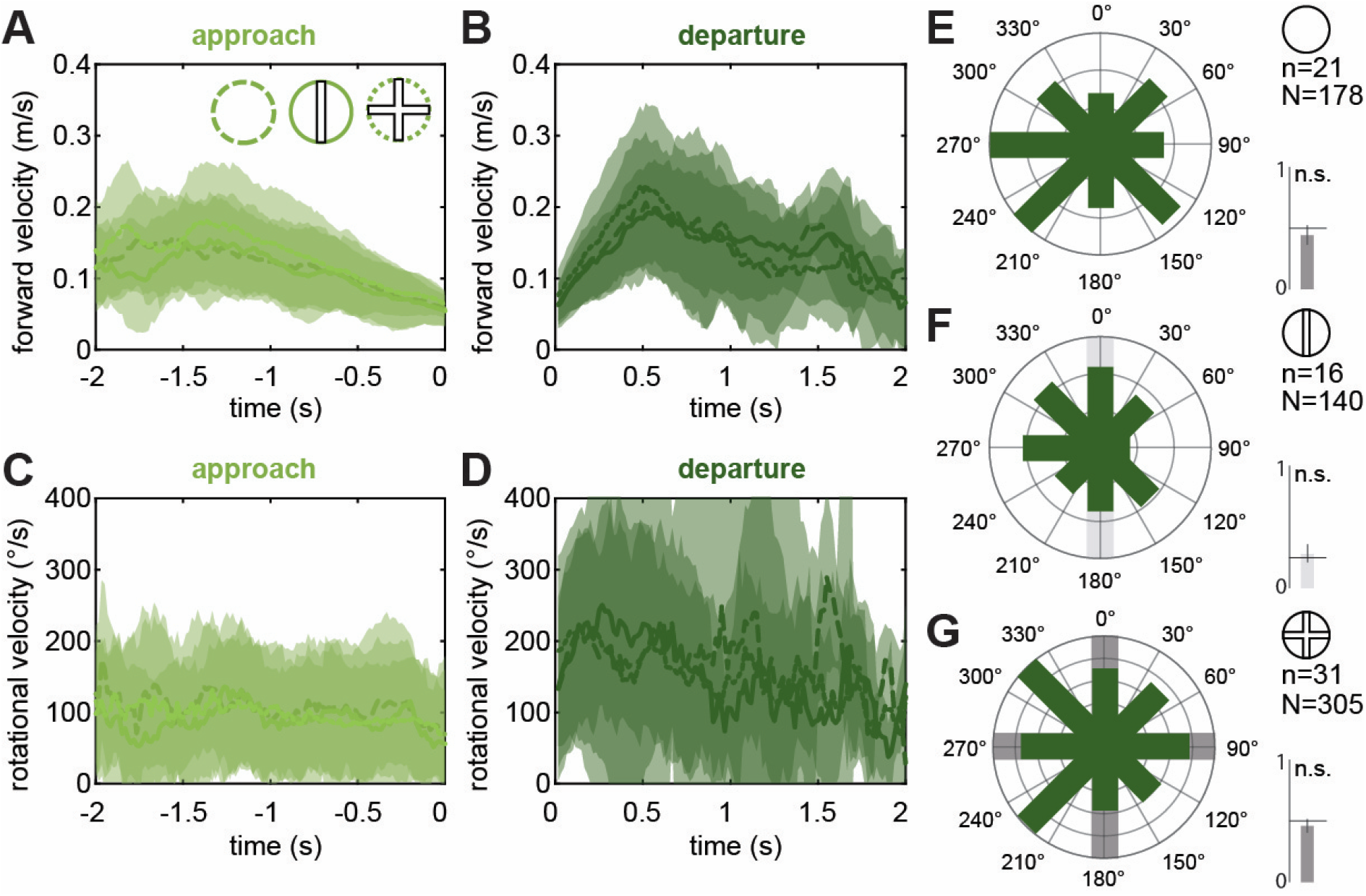
Approach and departure flight track features. **A-D** Forward velocity (**A**,**B**) and yaw rotation velocity (**C**,**D**) of the last 2 s before landing (*approach*, **A**,**C**) and after take-off (*departure*, **B**,**D**). Shown are mean and standard deviation for each pattern condition. Data is averaged over individual flight paths (N as in **E**,**F**,**G**), several of which were recorded for each individual bumblebee (n as in **E**,**F**,**G**). **E**,**F**,**G** Show circular histograms of the average heading angle of each bumblebee’s body axis (defined by the head and thorax position) in the first 500 ms of *departure* flights for the three pattern conditions. All data from conditions with patterns was re-orientated so that the patterns aligned with the cardinal axes (the *line* pattern always aligns with the Y-axis), while the *no* pattern condition remained in the original configuration. Thus, in the *no* pattern condition the cardinal axes represent the axes of the video camera’s field-of-view. The heading angles were binned into 8 sectors, of which 2 contain the pattern directions for the *line* pattern (0 and 180°, **F**) and 4 contain the pattern directions for the *cross* pattern (0, 90, 180 and 270°, **G**), as indicated by the grey shades. The statistical results of a generalised linear mixed-effects model (see Methods, Table S9, 10) implementing a binomial comparison of heading angles that fall into pattern sectors vs no-pattern sectors are shown as bar graphs next to each histogram. These graphs depict the mean and confidence intervals of heading angles that fell into pattern sectors, relative to the probability of landing in these sectors randomly.

**Fig. S3.**
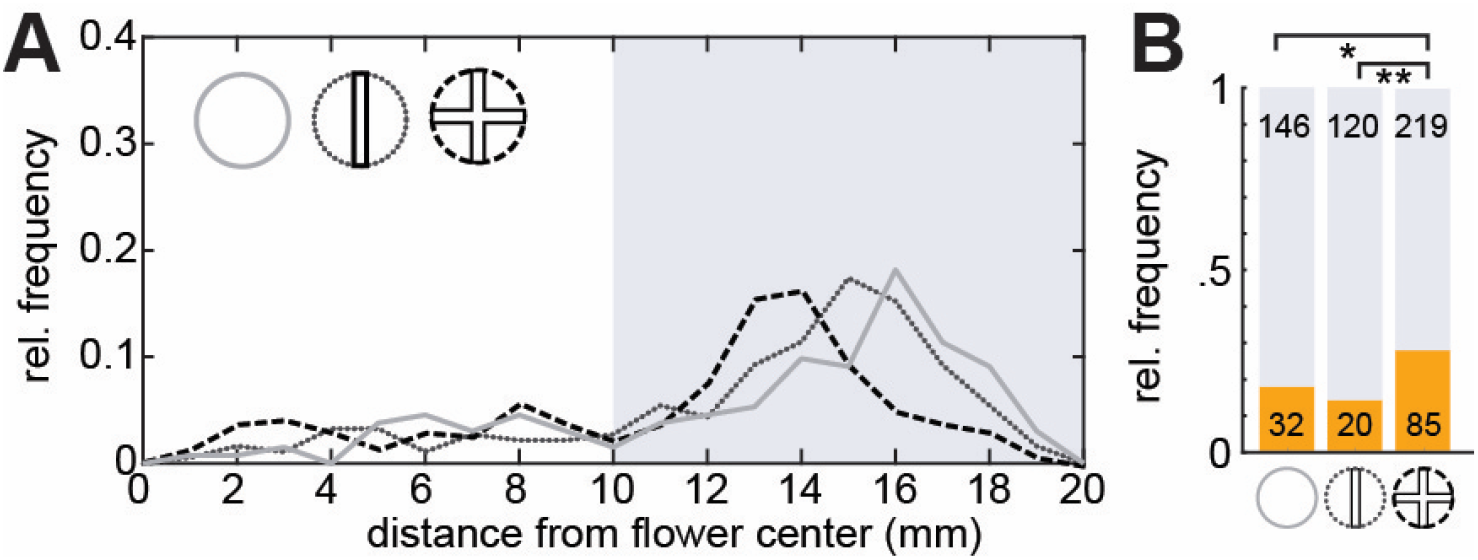
Landing position eccentricity for no, line and cross pattern conditions. **A** The relative frequency of landings for the three pattern types with respect to the distance from the flower centre, binned at 1 mm intervals. The shaded background highlights two groups for further analysis in **B**: landings closer than 10 mm to the centre and landings further than 10 mm. **B** The number of landings in the inner 10 mm (orange) and outer 10 mm (grey) of the flower was compared for all pattern types (Fisher’s exact test, • = 0.05, * < 0.05, ** < 0.01, *** < 0.001).

**Fig. S4.**
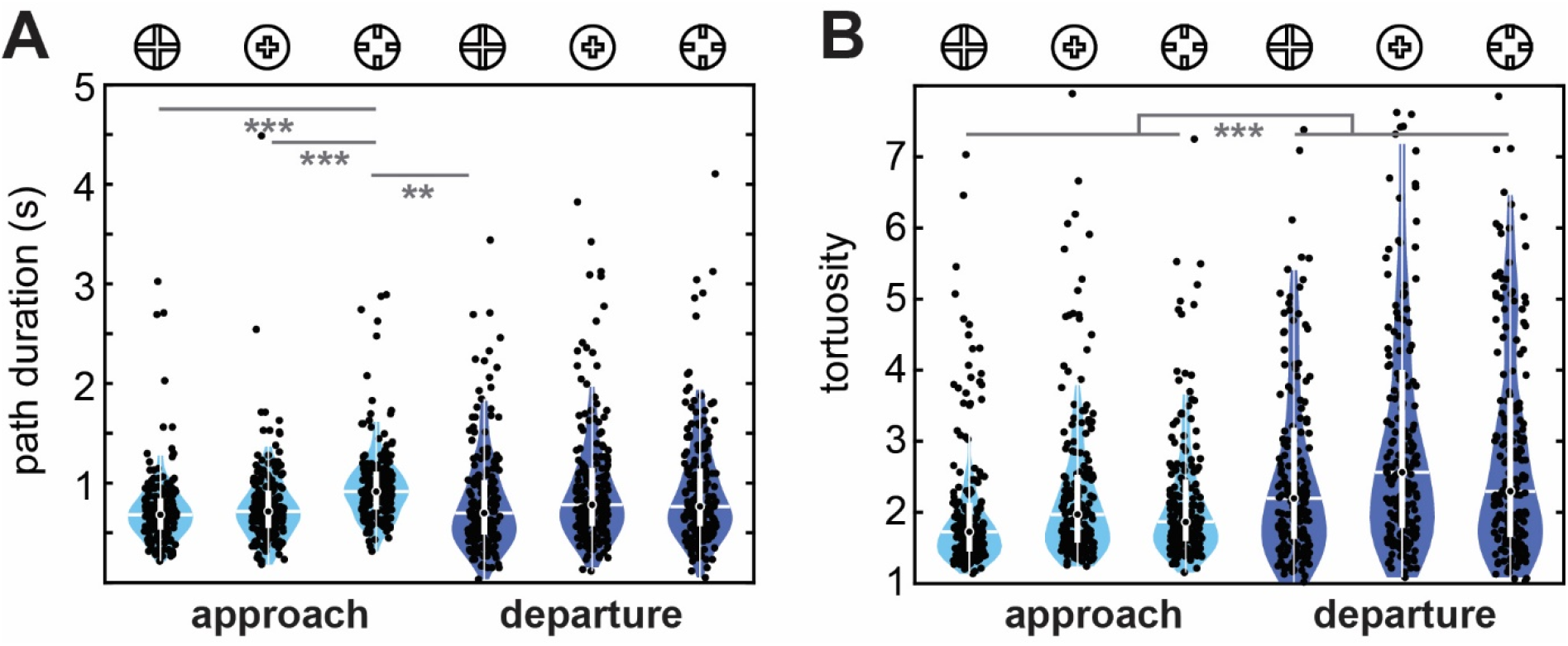
Walking paths on the flower with the cross patterns of varying reach. **A** Path duration, and **B** tortuosity of the *approach* and *departure* walks, respectively, in the *cross, inner cross* and *outer cross* conditions. Dots show data from individual flight paths (N: 200 in each condition), including multiple trials for individual bumblebees (n: 20 in each condition). The statistical results of a generalised linear mixed-effects model are indicated (see Methods, Tables S7, S8). Only significant results of the pairwise comparisons of all conditions (pattern type, flight type) are shown.

### Statistical results

**Table S1.** Statistical results of the generalised linear mixed-effects model for **total handling time**. *Lmm.allbg* is the model with *patternColour* (G = yellow, B = blue) as a factor: *track parameter* ~ *patternType × patternColour* + (1 + *patternType* × *patternColour* | *animalID*) and *lmm.all* without it: *track parameter ~ patternType* + (1 + *patternType* | *animalID*). Both models contain *patternType* (no, line, cross) as a factor.

| Model selection: Likelihood Ratio Test |  |  |  |  |  |  |  |  |
| --- | --- | --- | --- | --- | --- | --- | --- | --- |
|  | npar | AIC | BIC | logLik | deviance | Chisq | Df | Pr(>Chisq) |
| Imm.all | 15 | 652.43 | 718.92 | -311.21 | 622.43 |  |  |  |
| Imm.allbg | 28 | 630.7 | 754.82 | -287.35 | 574.7 | 47.728 | 13 | p < 0.001 |
| Multiple Comparisons of Means: Tukey Contrasts |  |  |  |  |  |  |  |  |
| Conditions |  | Estimate | Std. Error | z value |  | p value |  |  |
| no | line | -0.011 | 0.077 | -0.148 |  | 0.988 |  |  |
| no | cross | -0.249 | 0.079 | -3.162 |  | 0.004 |  |  |
| line | cross | -0.237 | 0.097 | -2.444 |  | 0.038 |  |  |
| G | B | -0.043 | 0.066 | -0.655 |  | 0.512 |  |  |
|  |  | Estimate | Std. Error | t ratio |  | p value |  |  |
| B/no | B/line | 0.011 | 0.085 | 0.134 |  | 1.000 |  |  |
| B/no | B/cross | 0.249 | 0.085 | 2.929 |  | 0.069 |  |  |
| B/no | G/no | 0.043 | 0.073 | 0.588 |  | 0.990 |  |  |
| B/no | G/line | 0.263 | 0.099 | 2.658 |  | 0.150 |  |  |
| B/no | G/cross | 0.200 | 0.080 | 2.490 |  | 0.178 |  |  |
| B/line | B/cross | 0.237 | 0.109 | 2.183 |  | 0.296 |  |  |
| B/line | G/no | 0.032 | 0.099 | 0.320 |  | 0.999 |  |  |
| B/line | G/line | 0.252 | 0.127 | 1.984 |  | 0.394 |  |  |
| B/line | G/cross | 0.189 | 0.112 | 1.680 |  | 0.562 |  |  |
| B/cross | G/no | -0.206 | 0.075 | -2.746 |  | 0.105 |  |  |
| B/cross | G/line | 0.014 | 0.116 | 0.123 |  | 1.000 |  |  |
| B/cross | G/cross | -0.049 | 0.067 | -0.727 |  | 0.977 |  |  |
| G/no | G/line | 0.220 | 0.090 | 2.458 |  | 0.202 |  |  |
| G/no | G/cross | 0.157 | 0.077 | 2.050 |  | 0.350 |  |  |
| G/line | G/cross | -0.063 | 0.105 | -0.603 |  | 0.989 |  |  |

**Table S2.**
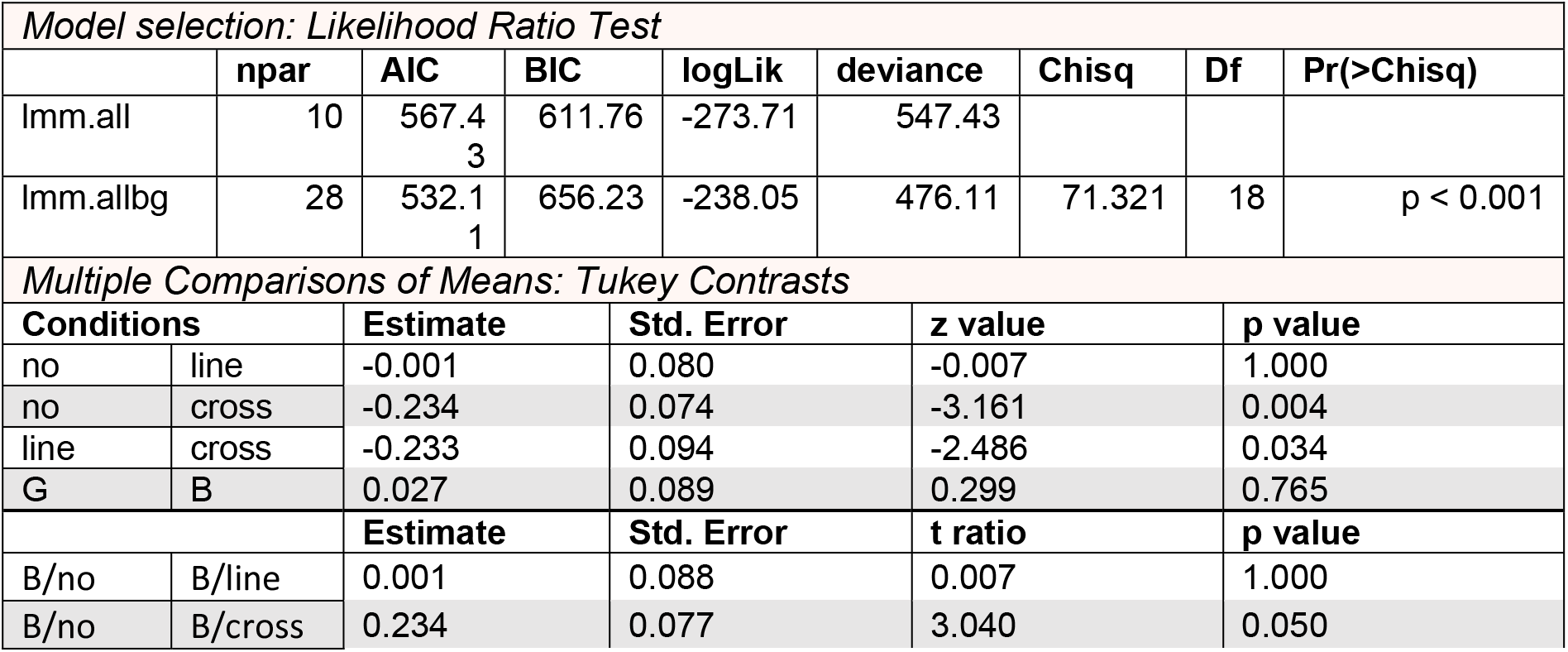
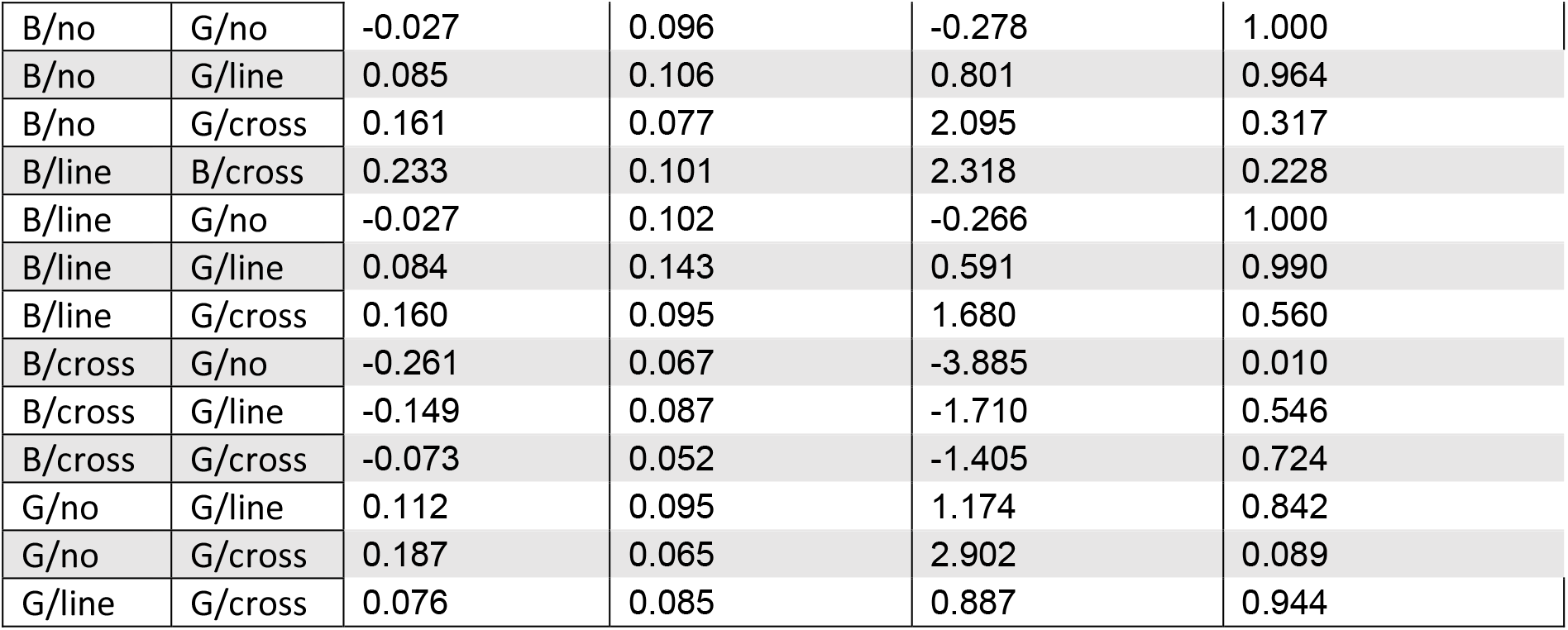
Statistical results of the generalised linear mixed-effects model for **approach duration**. *Lmm.allbg* is the model with *patternColour* (G = yellow, B = blue) as a factor, and *lmm.all* without it. Both models contain *patternType* (no, line, cross) as a factor.

| Model selection: Likelihood Ratio Test |  |  |  |  |  |  |  |  |
| --- | --- | --- | --- | --- | --- | --- | --- | --- |
|  | npar | AIC | BIC | logLik | deviance | Chisq | Df | Pr(>Chisq) |
| Imm.all | 10 | 567.43 | 611.76 | -273.71 | 547.43 |  |  |  |
| Imm.allbg | 28 | 532.11 | 656.23 | -238.05 | 476.11 | 71.321 | 18 | p < 0.001 |
| Multiple Comparisons of Means: Tukey Contrasts |  |  |  |  |  |  |  |  |
| Conditions |  | Estimate | Std. Error | z value |  | p value |  |  |
| no | line | -0.001 | 0.080 | -0.007 |  | 1.000 |  |  |
| no | cross | -0.234 | 0.074 | -3.161 |  | 0.004 |  |  |
| line | cross | -0.233 | 0.094 | -2.486 |  | 0.034 |  |  |
| G | B | 0.027 | 0.089 | 0.299 |  | 0.765 |  |  |
|  |  | Estimate | Std. Error | t ratio |  | p value |  |  |
| B/no | B/line | 0.001 | 0.088 | 0.007 |  | 1.000 |  |  |
| B/no | B/cross | 0.234 | 0.077 | 3.040 |  | 0.050 |  |  |

|  |  |  |  |  |  |
| --- | --- | --- | --- | --- | --- |
| B/no | G/no | -0.027 | 0.096 | -0.278 | 1.000 |
| B/no | G/line | 0.085 | 0.106 | 0.801 | 0.964 |
| B/no | G/cross | 0.161 | 0.077 | 2.095 | 0.317 |
| B/line | B/cross | 0.233 | 0.101 | 2.318 | 0.228 |
| B/line | G/no | -0.027 | 0.102 | -0.266 | 1.000 |
| B/line | G/line | 0.084 | 0.143 | 0.591 | 0.990 |
| B/line | G/cross | 0.160 | 0.095 | 1.680 | 0.560 |
| B/cross | G/no | -0.261 | 0.067 | -3.885 | 0.010 |
| B/cross | G/line | -0.149 | 0.087 | -1.710 | 0.546 |
| B/cross | G/cross | -0.073 | 0.052 | -1.405 | 0.724 |
| G/no | G/line | 0.112 | 0.095 | 1.174 | 0.842 |
| G/no | G/cross | 0.187 | 0.065 | 2.902 | 0.089 |
| G/line | G/cross | 0.076 | 0.085 | 0.887 | 0.944 |

**Table S3.** Statistical results of the generalised linear mixed-effects model for **forward velocity**. *Lmm.allbg* is the model with *patternColour* (G = yellow, B = blue) as a factor: *track parameter* ~ *foragingPhase × patternType × patternColour* + (1 + *foragingPhase × patternType × patternColour* | *animalID*) and *lmm.all* without it: *track parameter ~ foragingPhase × patternType* + (1 + *foragingPhase × patternType* | *animalID*). Both models contain *patternType* (no, line, cross) and *foragingPhase* (a = approach, d = departure) as a factor.

| Model selection: Likelihood Ratio Test |  |  |  |  |  |  |  |  |
| --- | --- | --- | --- | --- | --- | --- | --- | --- |
|  | npar | AIC | BIC | logLik | deviance | Chisq | Df | Pr(>Chisq) |
| Imm.all | 28 | 1175.5 | 1318.6 | -559.75 | 1119.5 |  |  |  |
| Imm.allbg | 91 | 1221.8 | 1687 | -519.92 | 1039.8 | 79.656 | 63 | 0.07659 |
| Multiple Comparisons of Means: Tukey Contrasts |  |  |  |  |  |  |  |  |
| Conditions |  | Estimate | Std. Error | z value | p value |  |  |  |
| no | line | -0.172 | 0.055 | -3.149 | 0.004 |  |  |  |
| no | cross | <0.001 | 0.052 | -0.003 | 1.000 |  |  |  |
| line | cross | 0.172 | 0.070 | 2.470 | 0.035 |  |  |  |
| a | d | 0.417 | 0.089 | 4.679 | <0.001 |  |  |  |
|  |  | Estimate | Std. Error | t ratio | p value |  |  |  |
| a/no | d/no | -0.417 | 0.095 | -4.380 | 0.003 |  |  |  |
| a/no | a/line | 0.172 | 0.061 | 2.831 | 0.104 |  |  |  |
| a/no | d/line | -0.340 | 0.073 | -4.656 | 0.002 |  |  |  |
| a/no | a/cross | <0.001 | 0.057 | 0.003 | 1.000 |  |  |  |
| a/no | d/cross | -0.502 | 0.077 | -6.547 | <0.001 |  |  |  |
| d/no | a/line | 0.589 | 0.113 | 5.223 | <0.001 |  |  |  |
| d/no | d/line | 0.078 | 0.060 | 1.292 | 0.786 |  |  |  |
| d/no | a/cross | 0.418 | 0.080 | 5.233 | <0.001 |  |  |  |
| d/no | d/cross | -0.085 | 0.084 | -1.014 | 0.909 |  |  |  |
| a/line | d/line | -0.512 | 0.087 | -5.874 | <0.001 |  |  |  |
| a/line | a/cross | -0.172 | 0.076 | -2.255 | 0.257 |  |  |  |
| a/line | d/cross | -0.674 | 0.097 | -6.990 | <0.001 |  |  |  |
| d/line | a/cross | 0.340 | 0.058 | 5.827 | <0.001 |  |  |  |
| d/line | d/cross | -0.162 | 0.071 | -2.281 | 0.238 |  |  |  |
| a/cross | d/cross | -0.503 | 0.051 | -9.948 | <0.001 |  |  |  |

**Table S4.** Statistical results of the generalised linear mixed-effects model for **rotational velocity**. *Lmm.allbg* is the model with *patternColour* (G = yellow, B = blue) as a factor, and *lmm.all* without it. Both models contain *patternType* (no, line, cross) and *foragingPhase* (a = approach, d = departure) as a factor.

| Model selection: Likelihood Ratio Test |  |  |  |  |  |  |  |  |
| --- | --- | --- | --- | --- | --- | --- | --- | --- |
|  | npars | AIC | BIC | logLik | deviance | Chisq | Df | Pr(>Chisq) |
| lmm.all | 28 | 2182.4 | 2325.5 | -1063.2 | 2126.4 |  |  |  |
| lmm.allbg | 91 | 2259.2 | 2724.3 | -1038.6 | 2077.2 | 49.208 | 63 | 0.8982 |
| Multiple Comparisons of Means: Tukey Contrasts |  |  |  |  |  |  |  |  |
| Conditions |  | Estimate | Std. Error | z value | p value |  |  |  |
| no | line | -0.176 | 0.066 | -2.655 | 0.022 |  |  |  |
| no | cross | -0.113 | 0.065 | -1.727 | 0.194 |  |  |  |
| line | cross | 0.063 | 0.074 | 0.846 | 0.673 |  |  |  |
| a | d | 0.562 | 0.064 | 8.728 | <0.001 |  |  |  |
|  |  | Estimate | Std. Error | t ratio | p value |  |  |  |
| a/no | d/no | -0.562 | 0.070 | -8.008 | <0.001 |  |  |  |
| a/no | a/line | 0.176 | 0.074 | 2.366 | 0.227 |  |  |  |
| a/no | d/line | -0.585 | 0.087 | -6.757 | <0.001 |  |  |  |
| a/no | a/cross | 0.113 | 0.070 | 1.608 | 0.601 |  |  |  |
| a/no | d/cross | -0.551 | 0.080 | -6.920 | <0.001 |  |  |  |
| d/no | a/line | 0.738 | 0.081 | 9.087 | <0.001 |  |  |  |
| d/no | d/line | -0.023 | 0.082 | -0.282 | 1.000 |  |  |  |
| d/no | a/cross | 0.675 | 0.073 | 9.270 | <0.001 |  |  |  |
| d/no | d/cross | 0.011 | 0.079 | 0.137 | 1.000 |  |  |  |
| a/line | d/line | -0.761 | 0.099 | -7.724 | <0.001 |  |  |  |
| a/line | a/cross | -0.063 | 0.082 | -0.771 | 0.970 |  |  |  |
| a/line | d/cross | -0.727 | 0.093 | -7.861 | <0.001 |  |  |  |
| d/line | a/cross | 0.698 | 0.079 | 8.824 | <0.001 |  |  |  |
| d/line | d/cross | 0.034 | 0.082 | 0.416 | 0.998 |  |  |  |
| a/cross | d/cross | -0.664 | 0.056 | -11.973 | <0.001 |  |  |  |

**Table S5.**
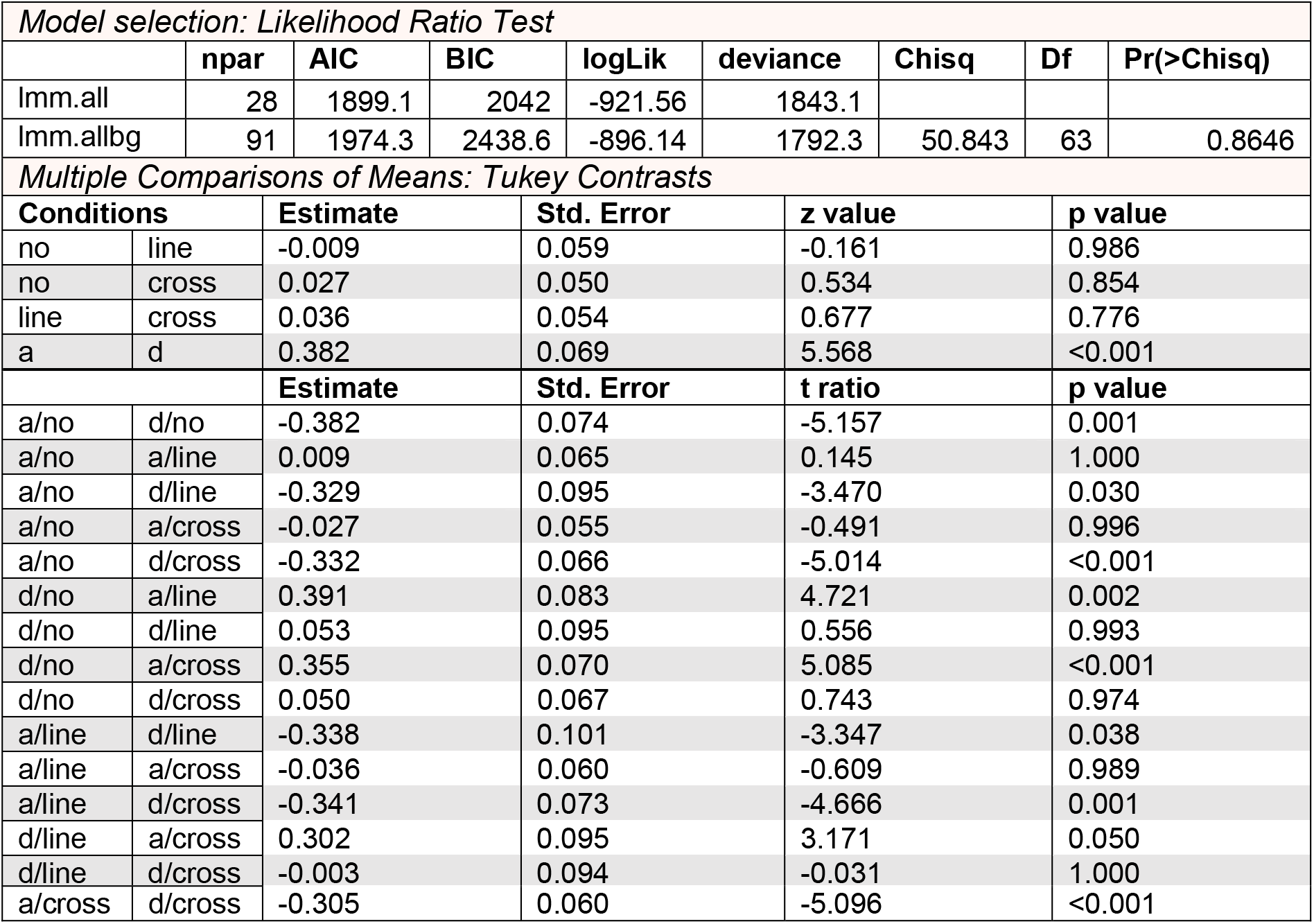
Statistical results of the generalised linear mixed-effects model for **walking path tortuosity**. *Lmm.allbg* is the model with *patternColour* (G = yellow, B = blue) as a factor, and *lmm.all* without it. Both models contain *patternType* (no, line, cross) and *foragingPhase* (a = approach, d = departure) as a factor.

| Model selection: Likelihood Ratio Test |  |  |  |  |  |  |  |  |
| --- | --- | --- | --- | --- | --- | --- | --- | --- |
|  | npars | AIC | BIC | logLik | deviance | Chisq | Df | Pr(>Chisq) |
| lmm.all | 28 | 1899.1 | 2042 | -921.56 | 1843.1 |  |  |  |
| lmm.allbg | 91 | 1974.3 | 2438.6 | -896.14 | 1792.3 | 50.843 | 63 | 0.8646 |
| Multiple Comparisons of Means: Tukey Contrasts |  |  |  |  |  |  |  |  |
| Conditions |  | Estimate | Std. Error | z value | p value |  |  |  |
| no | line | -0.009 | 0.059 | -0.161 | 0.986 |  |  |  |
| no | cross | 0.027 | 0.050 | 0.534 | 0.854 |  |  |  |
| line | cross | 0.036 | 0.054 | 0.677 | 0.776 |  |  |  |
| a | d | 0.382 | 0.069 | 5.568 | <0.001 |  |  |  |
|  |  | Estimate | Std. Error | t ratio | p value |  |  |  |
| a/no | d/no | -0.382 | 0.074 | -5.157 | 0.001 |  |  |  |
| a/no | a/line | 0.009 | 0.065 | 0.145 | 1.000 |  |  |  |
| a/no | d/line | -0.329 | 0.095 | -3.470 | 0.030 |  |  |  |
| a/no | a/cross | -0.027 | 0.055 | -0.491 | 0.996 |  |  |  |
| a/no | d/cross | -0.332 | 0.066 | -5.014 | <0.001 |  |  |  |
| d/no | a/line | 0.391 | 0.083 | 4.721 | 0.002 |  |  |  |
| d/no | d/line | 0.053 | 0.095 | 0.556 | 0.993 |  |  |  |
| d/no | a/cross | 0.355 | 0.070 | 5.085 | <0.001 |  |  |  |
| d/no | d/cross | 0.050 | 0.067 | 0.743 | 0.974 |  |  |  |
| a/line | d/line | -0.338 | 0.101 | -3.347 | 0.038 |  |  |  |
| a/line | a/cross | -0.036 | 0.060 | -0.609 | 0.989 |  |  |  |
| a/line | d/cross | -0.341 | 0.073 | -4.666 | 0.001 |  |  |  |
| d/line | a/cross | 0.302 | 0.095 | 3.171 | 0.050 |  |  |  |
| d/line | d/cross | -0.003 | 0.094 | -0.031 | 1.000 |  |  |  |

|  |  |  |  |  |  |
| --- | --- | --- | --- | --- | --- |
| a/cross | d/cross | -0.305 | 0.060 | -5.096 | <0.001 |

**Table S6.**
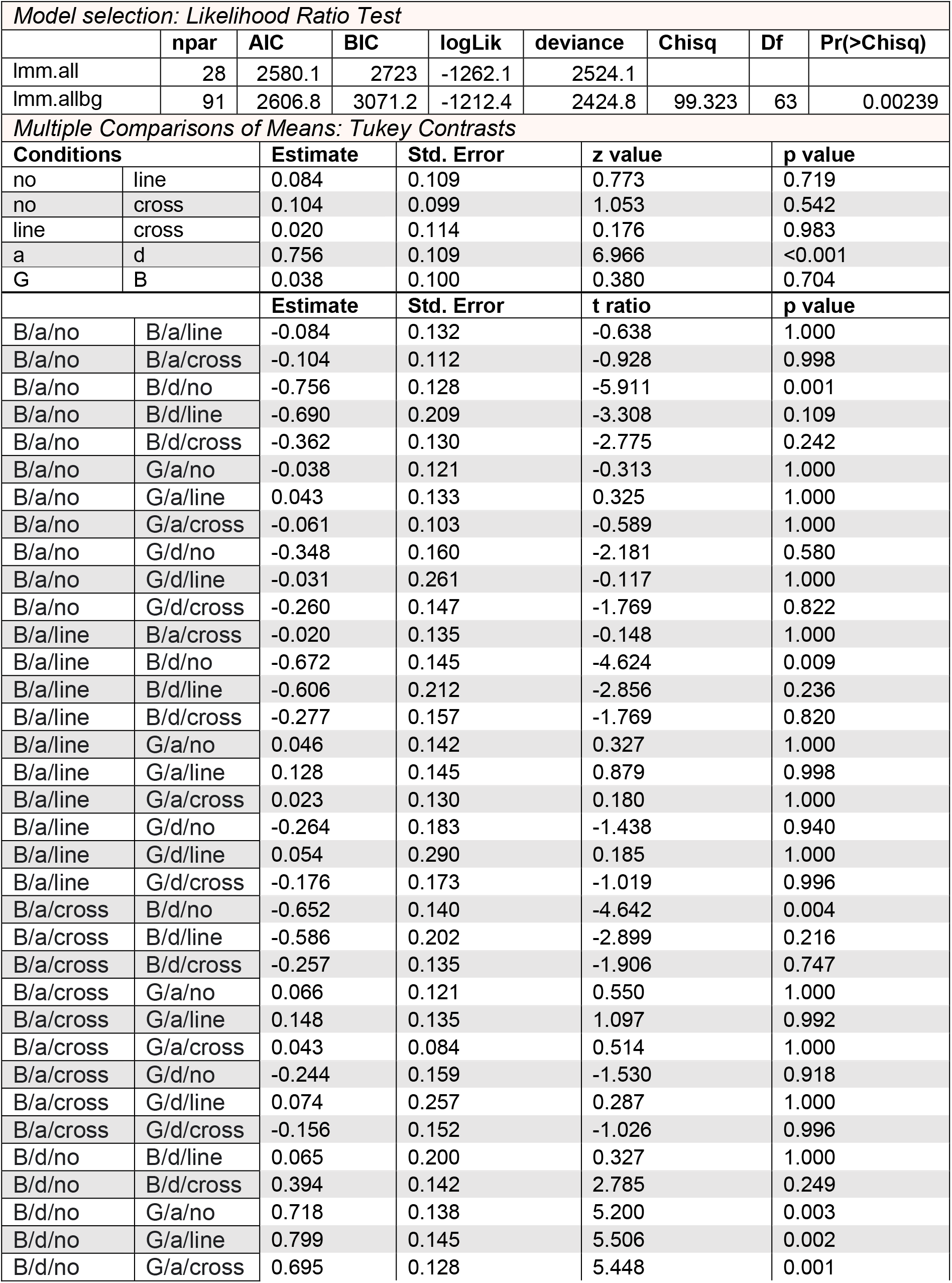
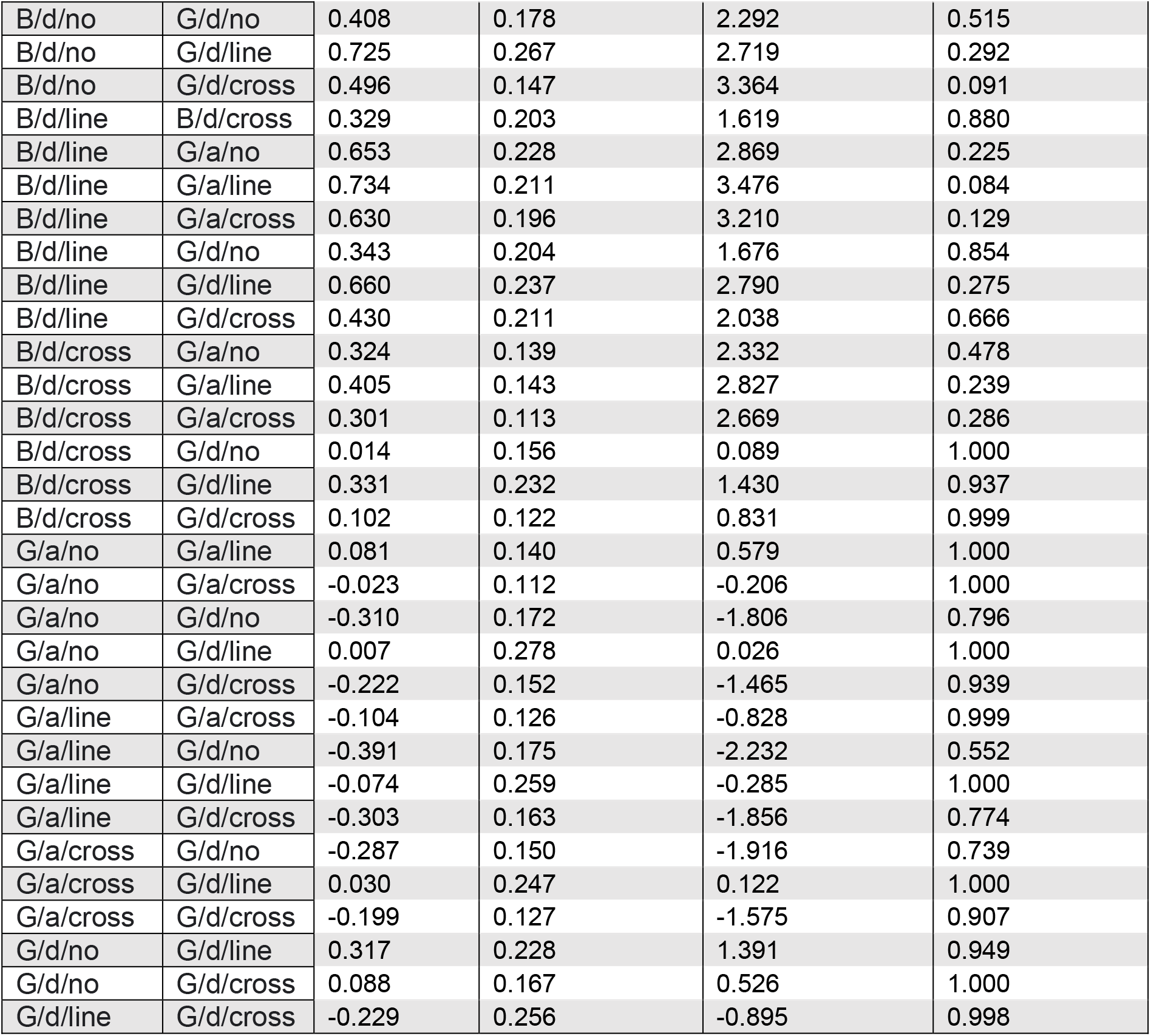
Statistical results of the generalised linear mixed-effects model for **walking path duration**. *Lmm.allbg* is the model with *patternColour* (G = yellow, B = blue) as a factor, and *lmm.all* without it. Both models contain *patternType* (no, line, cross) and *foragingPhase* (a = approach, d = departure) as a factor.

| Model selection: Likelihood Ratio Test |  |  |  |  |  |  |  |  |
| --- | --- | --- | --- | --- | --- | --- | --- | --- |
|  | npar | AIC | BIC | logLik | deviance | Chisq | Df | Pr(>Chisq) |
| lmm.all | 28 | 2580.1 | 2723 | -1262.1 | 2524.1 |  |  |  |
| lmm.allbg | 91 | 2606.8 | 3071.2 | -1212.4 | 2424.8 | 99.323 | 63 | 0.00239 |
| Multiple Comparisons of Means: Tukey Contrasts |  |  |  |  |  |  |  |  |
| Conditions |  | Estimate | Std. Error | z value | p value |  |  |  |
| no | line | 0.084 | 0.109 | 0.773 | 0.719 |  |  |  |
| no | cross | 0.104 | 0.099 | 1.053 | 0.542 |  |  |  |
| line | cross | 0.020 | 0.114 | 0.176 | 0.983 |  |  |  |
| a | d | 0.756 | 0.109 | 6.966 | <0.001 |  |  |  |
| G | B | 0.038 | 0.100 | 0.380 | 0.704 |  |  |  |
|  |  | Estimate | Std. Error | t ratio | p value |  |  |  |
| B/a/no | B/a/line | -0.084 | 0.132 | -0.638 | 1.000 |  |  |  |
| B/a/no | B/a/cross | -0.104 | 0.112 | -0.928 | 0.998 |  |  |  |
| B/a/no | B/d/no | -0.756 | 0.128 | -5.911 | 0.001 |  |  |  |
| B/a/no | B/d/line | -0.690 | 0.209 | -3.308 | 0.109 |  |  |  |
| B/a/no | B/d/cross | -0.362 | 0.130 | -2.775 | 0.242 |  |  |  |
| B/a/no | G/a/no | -0.038 | 0.121 | -0.313 | 1.000 |  |  |  |
| B/a/no | G/a/line | 0.043 | 0.133 | 0.325 | 1.000 |  |  |  |
| B/a/no | G/a/cross | -0.061 | 0.103 | -0.589 | 1.000 |  |  |  |
| B/a/no | G/d/no | -0.348 | 0.160 | -2.181 | 0.580 |  |  |  |
| B/a/no | G/d/line | -0.031 | 0.261 | -0.117 | 1.000 |  |  |  |
| B/a/no | G/d/cross | -0.260 | 0.147 | -1.769 | 0.822 |  |  |  |
| B/a/line | B/a/cross | -0.020 | 0.135 | -0.148 | 1.000 |  |  |  |
| B/a/line | B/d/no | -0.672 | 0.145 | -4.624 | 0.009 |  |  |  |
| B/a/line | B/d/line | -0.606 | 0.212 | -2.856 | 0.236 |  |  |  |
| B/a/line | B/d/cross | -0.277 | 0.157 | -1.769 | 0.820 |  |  |  |
| B/a/line | G/a/no | 0.046 | 0.142 | 0.327 | 1.000 |  |  |  |
| B/a/line | G/a/line | 0.128 | 0.145 | 0.879 | 0.998 |  |  |  |
| B/a/line | G/a/cross | 0.023 | 0.130 | 0.180 | 1.000 |  |  |  |
| B/a/line | G/d/no | -0.264 | 0.183 | -1.438 | 0.940 |  |  |  |
| B/a/line | G/d/line | 0.054 | 0.290 | 0.185 | 1.000 |  |  |  |
| B/a/line | G/d/cross | -0.176 | 0.173 | -1.019 | 0.996 |  |  |  |
| B/a/cross | B/d/no | -0.652 | 0.140 | -4.642 | 0.004 |  |  |  |
| B/a/cross | B/d/line | -0.586 | 0.202 | -2.899 | 0.216 |  |  |  |
| B/a/cross | B/d/cross | -0.257 | 0.135 | -1.906 | 0.747 |  |  |  |
| B/a/cross | G/a/no | 0.066 | 0.121 | 0.550 | 1.000 |  |  |  |
| B/a/cross | G/a/line | 0.148 | 0.135 | 1.097 | 0.992 |  |  |  |
| B/a/cross | G/a/cross | 0.043 | 0.084 | 0.514 | 1.000 |  |  |  |
| B/a/cross | G/d/no | -0.244 | 0.159 | -1.530 | 0.918 |  |  |  |
| B/a/cross | G/d/line | 0.074 | 0.257 | 0.287 | 1.000 |  |  |  |
| B/a/cross | G/d/cross | -0.156 | 0.152 | -1.026 | 0.996 |  |  |  |
| B/d/no | B/d/line | 0.065 | 0.200 | 0.327 | 1.000 |  |  |  |
| B/d/no | B/d/cross | 0.394 | 0.142 | 2.785 | 0.249 |  |  |  |
| B/d/no | G/a/no | 0.718 | 0.138 | 5.200 | 0.003 |  |  |  |
| B/d/no | G/a/line | 0.799 | 0.145 | 5.506 | 0.002 |  |  |  |
| B/d/no | G/a/cross | 0.695 | 0.128 | 5.448 | 0.001 |  |  |  |

|  |  |  |  |  |  |
| --- | --- | --- | --- | --- | --- |
| B/d/no | G/d/no | 0.408 | 0.178 | 2.292 | 0.515 |
| B/d/no | G/d/line | 0.725 | 0.267 | 2.719 | 0.292 |
| B/d/no | G/d/cross | 0.496 | 0.147 | 3.364 | 0.091 |
| B/d/line | B/d/cross | 0.329 | 0.203 | 1.619 | 0.880 |
| B/d/line | G/a/no | 0.653 | 0.228 | 2.869 | 0.225 |
| B/d/line | G/a/line | 0.734 | 0.211 | 3.476 | 0.084 |
| B/d/line | G/a/cross | 0.630 | 0.196 | 3.210 | 0.129 |
| B/d/line | G/d/no | 0.343 | 0.204 | 1.676 | 0.854 |
| B/d/line | G/d/line | 0.660 | 0.237 | 2.790 | 0.275 |
| B/d/line | G/d/cross | 0.430 | 0.211 | 2.038 | 0.666 |
| B/d/cross | G/a/no | 0.324 | 0.139 | 2.332 | 0.478 |
| B/d/cross | G/a/line | 0.405 | 0.143 | 2.827 | 0.239 |
| B/d/cross | G/a/cross | 0.301 | 0.113 | 2.669 | 0.286 |
| B/d/cross | G/d/no | 0.014 | 0.156 | 0.089 | 1.000 |
| B/d/cross | G/d/line | 0.331 | 0.232 | 1.430 | 0.937 |
| B/d/cross | G/d/cross | 0.102 | 0.122 | 0.831 | 0.999 |
| G/a/no | G/a/line | 0.081 | 0.140 | 0.579 | 1.000 |
| G/a/no | G/a/cross | -0.023 | 0.112 | -0.206 | 1.000 |
| G/a/no | G/d/no | -0.310 | 0.172 | -1.806 | 0.796 |
| G/a/no | G/d/line | 0.007 | 0.278 | 0.026 | 1.000 |
| G/a/no | G/d/cross | -0.222 | 0.152 | -1.465 | 0.939 |
| G/a/line | G/a/cross | -0.104 | 0.126 | -0.828 | 0.999 |
| G/a/line | G/d/no | -0.391 | 0.175 | -2.232 | 0.552 |
| G/a/line | G/d/line | -0.074 | 0.259 | -0.285 | 1.000 |
| G/a/line | G/d/cross | -0.303 | 0.163 | -1.856 | 0.774 |
| G/a/cross | G/d/no | -0.287 | 0.150 | -1.916 | 0.739 |
| G/a/cross | G/d/line | 0.030 | 0.247 | 0.122 | 1.000 |
| G/a/cross | G/d/cross | -0.199 | 0.127 | -1.575 | 0.907 |
| G/d/no | G/d/line | 0.317 | 0.228 | 1.391 | 0.949 |
| G/d/no | G/d/cross | 0.088 | 0.167 | 0.526 | 1.000 |
| G/d/line | G/d/cross | -0.229 | 0.256 | -0.895 | 0.998 |

**Table S7.**
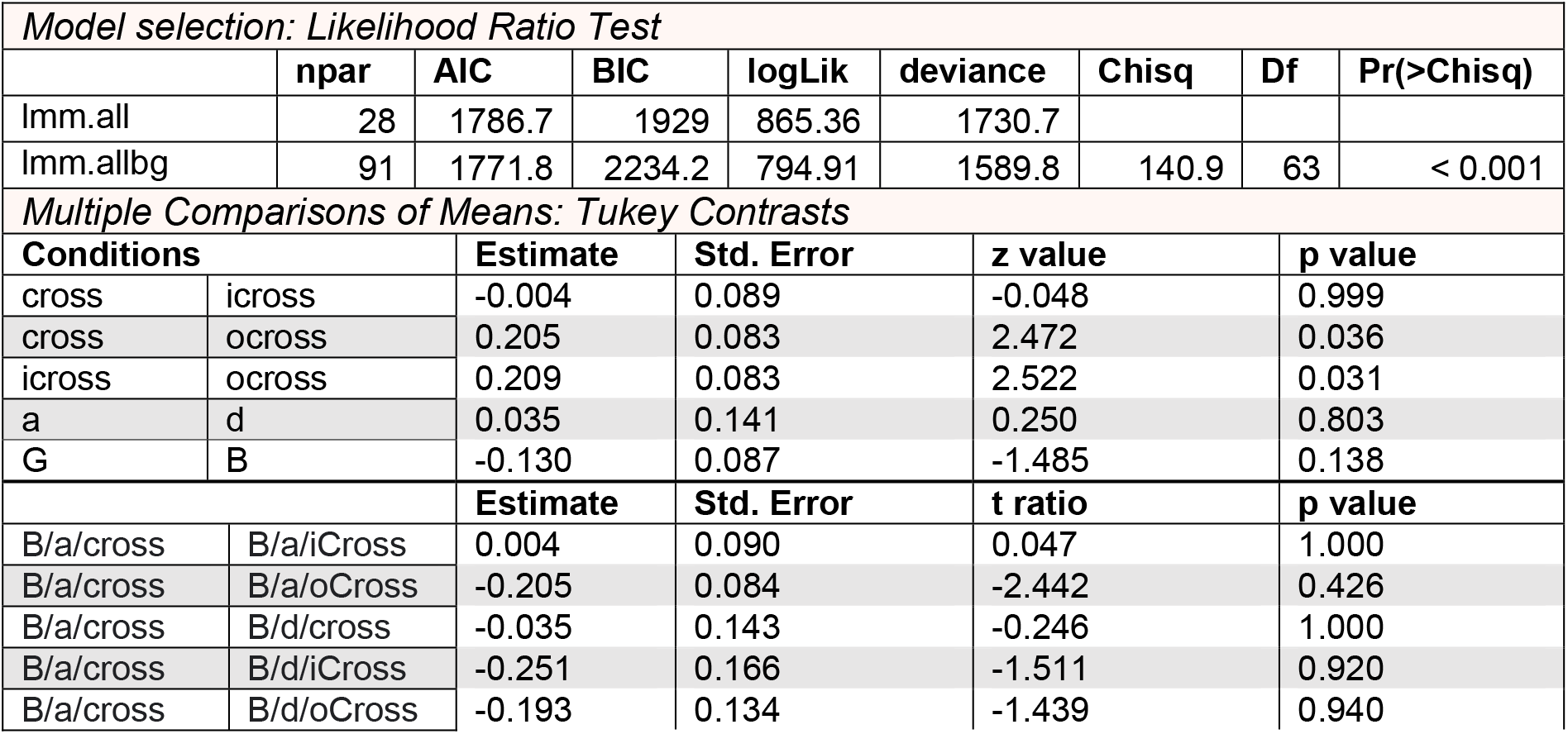
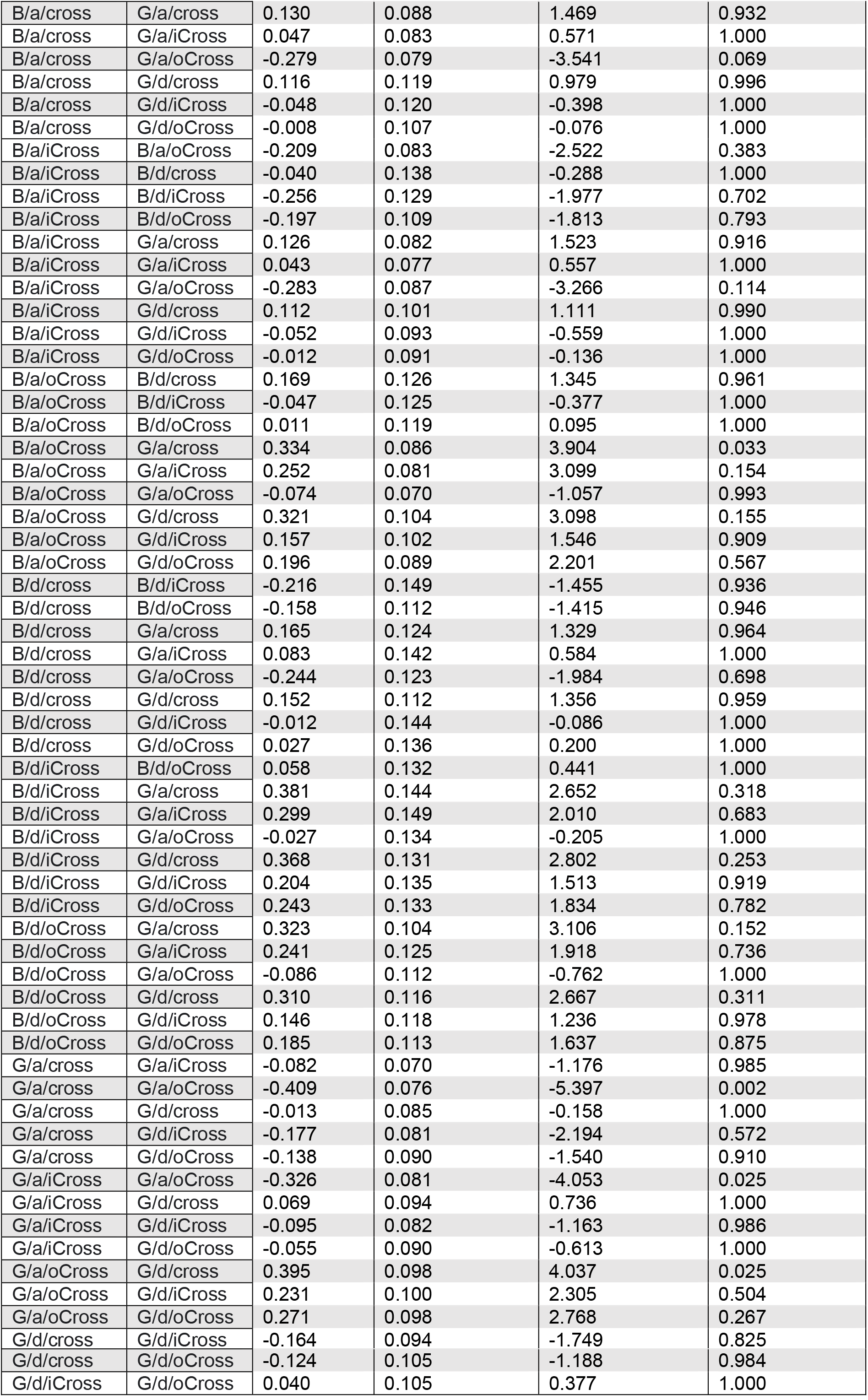
Statistical results of the generalised linear mixed-effects model for **walking path duration**. *Lmm.allbg* is the model with *patternColour* (G = yellow, B = blue) as a factor, and *lmm.all* without it. Both models contain *patternType* (no, line, cross) and *foragingPhase* (a = approach, d = departure) as a factor.

| Model selection: Likelihood Ratio Test |  |  |  |  |  |  |  |  |
| --- | --- | --- | --- | --- | --- | --- | --- | --- |
|  | npar | AIC | BIC | logLik | deviance | Chisq | Df | Pr(>Chisq) |
| Imm.all | 28 | 1786.7 | 1929 | 865.36 | 1730.7 |  |  |  |
| Imm.allbg | 91 | 1771.8 | 2234.2 | 794.91 | 1589.8 | 140.9 | 63 | < 0.001 |
| Multiple Comparisons of Means: Tukey Contrasts |  |  |  |  |  |  |  |  |
| Conditions |  | Estimate | Std. Error | z value |  | p value |  |  |
| cross | icross | -0.004 | 0.089 | -0.048 |  | 0.999 |  |  |
| cross | ocross | 0.205 | 0.083 | 2.472 |  | 0.036 |  |  |
| icross | ocross | 0.209 | 0.083 | 2.522 |  | 0.031 |  |  |
| a | d | 0.035 | 0.141 | 0.250 |  | 0.803 |  |  |
| G | B | -0.130 | 0.087 | -1.485 |  | 0.138 |  |  |
|  |  | Estimate | Std. Error | t ratio |  | p value |  |  |
| B/a/cross | B/a/iCross | 0.004 | 0.090 | 0.047 |  | 1.000 |  |  |
| B/a/cross | B/a/oCross | -0.205 | 0.084 | -2.442 |  | 0.426 |  |  |
| B/a/cross | B/d/cross | -0.035 | 0.143 | -0.246 |  | 1.000 |  |  |
| B/a/cross | B/d/iCross | -0.251 | 0.166 | -1.511 |  | 0.920 |  |  |
| B/a/cross | B/d/oCross | -0.193 | 0.134 | -1.439 |  | 0.940 |  |  |

|  |  |  |  |  |  |
| --- | --- | --- | --- | --- | --- |
| B/a/cross | G/a/cross | 0.130 | 0.088 | 1.469 | 0.932 |
| B/a/cross | G/a/iCross | 0.047 | 0.083 | 0.571 | 1.000 |
| B/a/cross | G/a/oCross | -0.279 | 0.079 | -3.541 | 0.069 |
| B/a/cross | G/d/cross | 0.116 | 0.119 | 0.979 | 0.996 |
| B/a/cross | G/d/iCross | -0.048 | 0.120 | -0.398 | 1.000 |
| B/a/cross | G/d/oCross | -0.008 | 0.107 | -0.076 | 1.000 |
| B/a/iCross | B/a/oCross | -0.209 | 0.083 | -2.522 | 0.383 |
| B/a/iCross | B/d/cross | -0.040 | 0.138 | -0.288 | 1.000 |
| B/a/iCross | B/d/iCross | -0.256 | 0.129 | -1.977 | 0.702 |
| B/a/iCross | B/d/oCross | -0.197 | 0.109 | -1.813 | 0.793 |
| B/a/iCross | G/a/cross | 0.126 | 0.082 | 1.523 | 0.916 |
| B/a/iCross | G/a/iCross | 0.043 | 0.077 | 0.557 | 1.000 |
| B/a/iCross | G/a/oCross | -0.283 | 0.087 | -3.266 | 0.114 |
| B/a/iCross | G/d/cross | 0.112 | 0.101 | 1.111 | 0.990 |
| B/a/iCross | G/d/iCross | -0.052 | 0.093 | -0.559 | 1.000 |
| B/a/iCross | G/d/oCross | -0.012 | 0.091 | -0.136 | 1.000 |
| B/a/oCross | B/d/cross | 0.169 | 0.126 | 1.345 | 0.961 |
| B/a/oCross | B/d/iCross | -0.047 | 0.125 | -0.377 | 1.000 |
| B/a/oCross | B/d/oCross | 0.011 | 0.119 | 0.095 | 1.000 |
| B/a/oCross | G/a/cross | 0.334 | 0.086 | 3.904 | 0.033 |
| B/a/oCross | G/a/iCross | 0.252 | 0.081 | 3.099 | 0.154 |
| B/a/oCross | G/a/oCross | -0.074 | 0.070 | -1.057 | 0.993 |
| B/a/oCross | G/d/cross | 0.321 | 0.104 | 3.098 | 0.155 |
| B/a/oCross | G/d/iCross | 0.157 | 0.102 | 1.546 | 0.909 |
| B/a/oCross | G/d/oCross | 0.196 | 0.089 | 2.201 | 0.567 |
| B/d/cross | B/d/iCross | -0.216 | 0.149 | -1.455 | 0.936 |
| B/d/cross | B/d/oCross | -0.158 | 0.112 | -1.415 | 0.946 |
| B/d/cross | G/a/cross | 0.165 | 0.124 | 1.329 | 0.964 |
| B/d/cross | G/a/iCross | 0.083 | 0.142 | 0.584 | 1.000 |
| B/d/cross | G/a/oCross | -0.244 | 0.123 | -1.984 | 0.698 |
| B/d/cross | G/d/cross | 0.152 | 0.112 | 1.356 | 0.959 |
| B/d/cross | G/d/iCross | -0.012 | 0.144 | -0.086 | 1.000 |
| B/d/cross | G/d/oCross | 0.027 | 0.136 | 0.200 | 1.000 |
| B/d/iCross | B/d/oCross | 0.058 | 0.132 | 0.441 | 1.000 |
| B/d/iCross | G/a/cross | 0.381 | 0.144 | 2.652 | 0.318 |
| B/d/iCross | G/a/iCross | 0.299 | 0.149 | 2.010 | 0.683 |
| B/d/iCross | G/a/oCross | -0.027 | 0.134 | -0.205 | 1.000 |
| B/d/iCross | G/d/cross | 0.368 | 0.131 | 2.802 | 0.253 |
| B/d/iCross | G/d/iCross | 0.204 | 0.135 | 1.513 | 0.919 |
| B/d/iCross | G/d/oCross | 0.243 | 0.133 | 1.834 | 0.782 |
| B/d/oCross | G/a/cross | 0.323 | 0.104 | 3.106 | 0.152 |
| B/d/oCross | G/a/iCross | 0.241 | 0.125 | 1.918 | 0.736 |
| B/d/oCross | G/a/oCross | -0.086 | 0.112 | -0.762 | 1.000 |
| B/d/oCross | G/d/cross | 0.310 | 0.116 | 2.667 | 0.311 |
| B/d/oCross | G/d/iCross | 0.146 | 0.118 | 1.236 | 0.978 |
| B/d/oCross | G/d/oCross | 0.185 | 0.113 | 1.637 | 0.875 |
| G/a/cross | G/a/iCross | -0.082 | 0.070 | -1.176 | 0.985 |
| G/a/cross | G/a/oCross | -0.409 | 0.076 | -5.397 | 0.002 |
| G/a/cross | G/d/cross | -0.013 | 0.085 | -0.158 | 1.000 |
| G/a/cross | G/d/iCross | -0.177 | 0.081 | -2.194 | 0.572 |
| G/a/cross | G/d/oCross | -0.138 | 0.090 | -1.540 | 0.910 |
| G/a/iCross | G/a/oCross | -0.326 | 0.081 | -4.053 | 0.025 |
| G/a/iCross | G/d/cross | 0.069 | 0.094 | 0.736 | 1.000 |
| G/a/iCross | G/d/iCross | -0.095 | 0.082 | -1.163 | 0.986 |
| G/a/iCross | G/d/oCross | -0.055 | 0.090 | -0.613 | 1.000 |
| G/a/oCross | G/d/cross | 0.395 | 0.098 | 4.037 | 0.025 |
| G/a/oCross | G/d/iCross | 0.231 | 0.100 | 2.305 | 0.504 |
| G/a/oCross | G/d/oCross | 0.271 | 0.098 | 2.768 | 0.267 |
| G/d/cross | G/d/iCross | -0.164 | 0.094 | -1.749 | 0.825 |
| G/d/cross | G/d/oCross | -0.124 | 0.105 | -1.188 | 0.984 |
| G/d/iCross | G/d/oCross | 0.040 | 0.105 | 0.377 | 1.000 |

**Table S8.** Statistical results of the generalised linear mixed-effects model for **walking path tortuosity**. *Lmm.allbg* is the model with *patternColour* (G = yellow, B = blue) as a factor, and *lmm.all* without it. Both models contain *patternType* (no, line, cross) and *foragingPhase* (a = approach, d = departure) as a factor.

| Model selection: Likelihood Ratio Test |  |  |  |  |  |  |  |  |
| --- | --- | --- | --- | --- | --- | --- | --- | --- |
|  | npar | AIC | BIC | logLik | deviance | Chisq | Df | Pr(>Chisq) |
| lmm.all | 28 | 1823.2 | 1965.5 | -883.58 | 1767.2 |  |  |  |
| lmm.allbg | 91 | 1890.6 | 2353 | -854.28 | 1708.6 | 58.601 | 63 | 0.6337 |
| Multiple Comparisons of Means: Tukey Contrasts |  |  |  |  |  |  |  |  |
| Conditions |  | Estimate | Std. Error | z value | p value |  |  |  |
| no | line | 0.114 | 0.050 | 2.262 | 0.061 |  |  |  |
| no | cross | 0.092 | 0.060 | 1.527 | 0.277 |  |  |  |
| line | cross | -0.022 | 0.061 | -0.364 | 0.929 |  |  |  |
| a | d | 0.223 | 0.063 | 3.512 | <0.001 |  |  |  |
|  |  | Estimate | Std. Error | t ratio | p value |  |  |  |
| a/cross | d/cross | -0.223 | 0.064 | -3.504 | 0.025 |  |  |  |
| a/cross | a/iCross | -0.114 | 0.051 | -2.258 | 0.259 |  |  |  |
| a/cross | d/iCross | -0.379 | 0.072 | -5.294 | 0.001 |  |  |  |
| a/cross | a/oCross | -0.092 | 0.060 | -1.525 | 0.654 |  |  |  |
| a/cross | d/oCross | -0.289 | 0.061 | -4.763 | 0.002 |  |  |  |
| d/cross | a/iCross | 0.109 | 0.061 | 1.783 | 0.499 |  |  |  |
| d/cross | d/iCross | -0.156 | 0.090 | -1.726 | 0.532 |  |  |  |
| d/cross | a/oCross | 0.131 | 0.066 | 1.986 | 0.386 |  |  |  |
| d/cross | d/oCross | -0.066 | 0.072 | -0.918 | 0.937 |  |  |  |
| a/iCross | d/iCross | -0.265 | 0.075 | -3.516 | 0.024 |  |  |  |
| a/iCross | a/oCross | 0.022 | 0.061 | 0.364 | 0.999 |  |  |  |
| a/iCross | d/oCross | -0.175 | 0.062 | -2.828 | 0.096 |  |  |  |
| d/iCross | a/oCross | 0.287 | 0.062 | 4.668 | 0.002 |  |  |  |
| d/iCross | d/oCross | 0.090 | 0.075 | 1.204 | 0.830 |  |  |  |
| a/oCross | d/oCross | -0.197 | 0.071 | -2.768 | 0.107 |  |  |  |

**Table S9.** Statistical results of the model comparison via likelihood ratio tests for each generalised linear mixed-effects models of relative probabilities of positions in pattern vs non-pattern sectors. *BG* is the model with *patternColour* as a factor: *sector choice* ~ *patternColour* + (1 + *patternColour* | *animalID*) and *noBG* without it: *sector choice ~ 1* + (1| *animalID*).

|  |  | model 1 | model 2 | deviance 1 | deviance 2 | X<sup>2</sup> | p |
| --- | --- | --- | --- | --- | --- | --- | --- |
| appr. heading | no | noBG | BG | 115.353 | 114.853 | 0.500 | 0.480 |
|  | line | noBG | BG | 89.586 | 89.510 | 0.076 | 0.783 |
|  | cross | noBG | BG | 176.415 | 176.312 | 0.103 | 0.748 |
| depr. heading | no | noBG | BG | 111.820 | 109.223 | 2.597 | 0.107 |
|  | line | noBG | BG | 71.759 | 71.758 | 0.002 | 0.968 |
|  | cross | noBG | BG | 180.008 | 179.938 | 0.070 | 0.792 |
| landing heading | no | noBG | BG | 102.058 | 101.477 | 0.581 | 0.446 |
|  | line | noBG | BG | 86.953 | 86.715 | 0.238 | 0.626 |
|  | cross | noBG | BG | 178.237 | 177.868 | 0.369 | 0.544 |
| landing position | no | noBG | BG | 98.084 | 97.517 | 0.567 | 0.452 |
|  | line | noBG | BG | 86.191 | 86.189 | 0.002 | 0.965 |
|  | cross | noBG | BG | 161.008 | 160.887 | 0.121 | 0.728 |
| landing position | cross | noBG | BG | 98.661 | 98.593 | 0.068 | 0.794 |
|  | icross | noBG | BG | 123.629 | 121.339 | 2.290 | 0.130 |
|  | ocross | noBG | BG | 112.548 | 112.364 | 0.184 | 0.668 |
| pattern landing | line | noBG | BG | 71.183 | 71.011 | 0.173 | 0.677 |
|  | cross | noBG | BG | 182.645 | 179.770 | 2.874 | 0.090 |

**Table S10.**
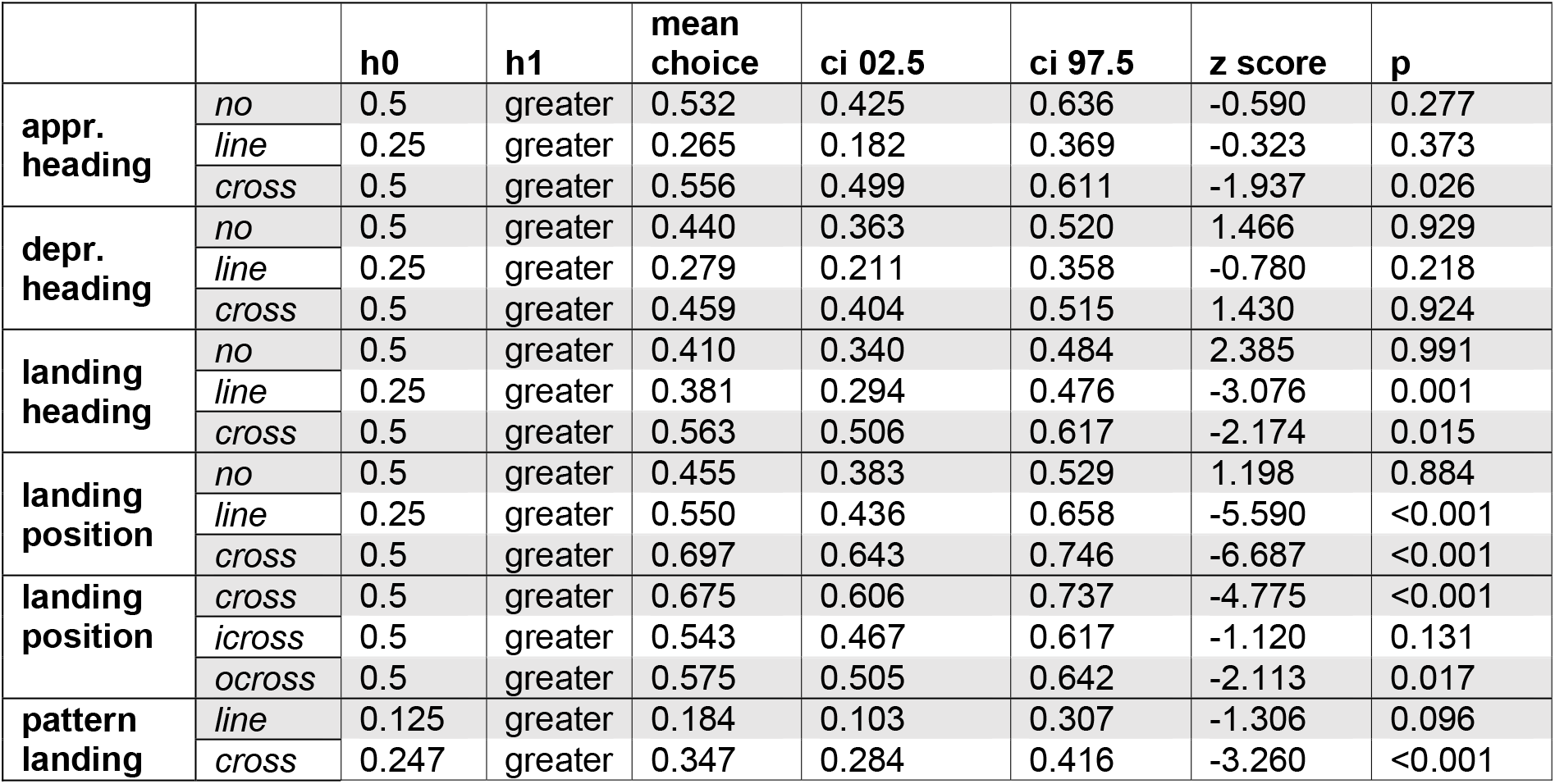
Statistical results of the generalised linear mixed-effects model to test distributions in pattern vs non-pattern sectors of the model *noBG* without the factor *patternColour*.

|  |  | h0 | h1 | mean choice | ci 02.5 | ci 97.5 | z score | p |
| --- | --- | --- | --- | --- | --- | --- | --- | --- |
| appr. heading | no | 0.5 | greater | 0.532 | 0.425 | 0.636 | -0.590 | 0.277 |
|  | line | 0.25 | greater | 0.265 | 0.182 | 0.369 | -0.323 | 0.373 |
|  | cross | 0.5 | greater | 0.556 | 0.499 | 0.611 | -1.937 | 0.026 |
| depr. heading | no | 0.5 | greater | 0.440 | 0.363 | 0.520 | 1.466 | 0.929 |
|  | line | 0.25 | greater | 0.279 | 0.211 | 0.358 | -0.780 | 0.218 |
|  | cross | 0.5 | greater | 0.459 | 0.404 | 0.515 | 1.430 | 0.924 |
| landing heading | no | 0.5 | greater | 0.410 | 0.340 | 0.484 | 2.385 | 0.991 |
|  | line | 0.25 | greater | 0.381 | 0.294 | 0.476 | -3.076 | 0.001 |
|  | cross | 0.5 | greater | 0.563 | 0.506 | 0.617 | -2.174 | 0.015 |
| landing position | no | 0.5 | greater | 0.455 | 0.383 | 0.529 | 1.198 | 0.884 |
|  | line | 0.25 | greater | 0.550 | 0.436 | 0.658 | -5.590 | <0.001 |
|  | cross | 0.5 | greater | 0.697 | 0.643 | 0.746 | -6.687 | <0.001 |
| landing position | cross | 0.5 | greater | 0.675 | 0.606 | 0.737 | -4.775 | <0.001 |
|  | icross | 0.5 | greater | 0.543 | 0.467 | 0.617 | -1.120 | 0.131 |
|  | ocross | 0.5 | greater | 0.575 | 0.505 | 0.642 | -2.113 | 0.017 |
| pattern landing | line | 0.125 | greater | 0.184 | 0.103 | 0.307 | -1.306 | 0.096 |
|  | cross | 0.247 | greater | 0.347 | 0.284 | 0.416 | -3.260 | <0.001 |

## References

Abrol, D. P. (2012), Pollination Biology, Springer Netherlands.

Baird, E., Boeddeker, N., Ibbotson, M. R. & Srinivasan, M. V. (2013), ‘A universal strategy for visually guided landing’, Proc. Natl. Acad. Sci. 110(46), 18686–18691.

Chang, J. J., Crall, J. D. & Combes, S. A. (2016), ‘Wind alters landing dynamics in bumblebees’, J. Exp. Biol. 219(18):2819–2822.

Chittka, L., Shmida, A., Troje, N. & Menzel, R. (1994), ‘Ultraviolet as a component of flower reflections, and the colour perception of Hymenoptera’, Vision Res. 34(11), 1489–1508.

Dafni, A. (1984), ‘Mimicry and deception in pollination.’, Annu. Rev. Ecol. Evol. Syst. 4, 259–278.

Dafni, A. & Giurfa, M. (1999), The functional ecology of floral guides in relation to insects behaviour and vision, in ‘Evolutionary Theory and Processes: Modern Perspectives’, Springer Netherlands, pp. 363–383.

Dafni, A. & Kevan, P. G. (1996), ‘Floral symmetry and nectar guides: ontogenetic constraints from floral development, colour pattern rules and functional significance’, Bot. J. Linn. Soc. 120(4), 371–377.

Dafni, A. & Kevan, P. G. (1997), ‘Flower size and shape: implications in pollination’, Isr. J. Plant. Sci. 45(2-3), 201–211.

Dafni, A., Lehrer, M. & Kevan, P. G. (1997), ‘Spatial flower parameters and insect spatial vision’, Biol. Rev. 72(2), 239–282.

de Ibarra, N. H., Langridge, K. V. & Vorobyev, M. (2015), ‘More than colour attraction: behavioural functions of flower patterns’, Curr. Opin. Insect Sci. 12, 64–70.

de Ibarra, N. H., M., G. & M., V. (2002), ‘Discrimination of coloured patterns by honeybees through chromatic and achromatic cues’, J. Comp. Physiol. A. 188, 503–512.

De Ibarra, N. H. & Vorobyev, M. (2009), ‘Flower patterns are adapted for detection by bees’, J. Comp. Physiol. A. 195(3), 319–323.

Dinkel, T. & Lunau, K. (2001), ‘How drone flies (*Eristalis tenax* l., Syrphidae, Diptera) use floral guides to locate food sources’, J. Insect Physiol. 47, 1111–1118.

Free, J. (1970), ‘Effect of flower shapes and nectar guides on the behaviour of foraging honeybees’, Behaviour 37(3-4), 269–285.

Garcia, J. E. & Dyer, A. G. (2021), ‘False colour photography reveals the complexity of flower signalling. a commentary on: ‘a bee’s eye view of remarkable floral colour patterns in the southwest Australian biodiversity hotspot revealed by false colour photography”, Annals Bot. 128(7), 10.1093/aob/mcab076.

Garcia, J. E., Greentree, A. D., Shrestha, M., Dorin, A. & Dyer, A. G. (2014), ‘Flower colours through the lens: Quantitative measurement with visible and ultraviolet digital photography’, PLoS ONE 9(5), e96646.

Goodale, E., Kim, E., Nabors, A., Henrichon, S. & Nieh, J. C. (2014), ‘The innate responses of bumble bees to flower patterns: separating the nectar guide from the nectary changes bee movements and search time.’, Naturwiss. 101, 523–526.

Goyret, J. (2010), ‘Look and touch: multimodal sensory control of flower inspection movements in the nocturnal hawkmoth *Manduca sexta*.’, J. Exp. Biol. 213(Pt 21), 3676–3682.

Goyret, J. & Kelber, A. (2012), ‘Chromatic signals control proboscis movements during hovering flight in the hummingbird hawkmoth *Macroglossum stellatarum*.’, PLoS One 7(4), e34629.

Hansen, D. M., Van der Niet, T. & Johnson, S. D. (2011), ‘Floral signposts: testing the significance of visual ‘nectar guides’ for pollinator behaviour and plant fitness’, Phil. Trans. R. Soc. Lond. B. 279(1729), 634–639.

Hedrick, T. L. (2008), ‘Software techniques for two- and three-dimensional kinematic measurements of biological and biomimetic systems.’, Bioinspir. Biomim. 3(3), 034001.

Heuschen, B., Gumbert, A. & Lunau, K. (2005), ‘A generalised mimicry system involving angiosperm flower colour, pollen and bumblebees’ innate colour preferences’, Plant. Syst. Evol. 252(3-4), 121–137.

Johnson, S. & Dafni, A. (1998), ‘Response of bee-flies to the shape and pattern of model flowers: implications for floral evolution in a mediterranean herb’, Funct. Ecol. 12(2), 289–297.

Kelber, A. (2002), ‘Pattern discrimination in a hawkmoth: Innate preferences, learning performance and ecology’, Proc. R. Soc. B. 269, 2573–2577.

Latty, T. & Trueblood, J. S. (2020), ‘How do insects choose flowers? a review of multi-attribute flower choice and decoy effects in flower-visiting insects’, J. Anim. Ecol. 89(12), 2750–2762.

Lawson, D. A., Whitney, H. M. & Rands, S. A. (2017), ‘Nectar discovery speeds and multimodal displays: assessing nectar search times in bees with radiating and non-radiating guides’, Evol. Ecol. 31(6), 899–912.

Lehrer, M. & Srinivasan, M. (1993), ‘Object detection by honeybees: Why do they land on edges?’, J. Comp. Physiol. A. 173(1), 23–32.

Leonard, A. S. & Papaj, D. R. (2011), ‘X marks the spot: The possible benefits of nectar guides to bees and plants’, Funct. Ecol. 25(6), 1293–1301.

Lunau, K., Scaccabarozzi, D., Willing, L. & Dixon, K. (2021), ‘A bee’s eye view of remarkable floral colour patterns in the southwest australian biodiversity hotspot revealed by false colour photography’, Annals Bot. 128(7), 821–824.

Lunau, K., Unseld, K. & Wolter, F. (2009), ‘Visual detection of diminutive floral guides in the bumblebee bombus terrestris and in the honeybee apis mellifera’, J. Comp. Physiol. A. 195, 1121–1130.

Maimon, G., Straw, A. D. & Dickinson, M. H. (2008), ‘A simple vision-based algorithm for decision making in flying drosophila’, Curr. Biol. 18(6), 464–470.

Makino, T. T. (2008), ‘Bumble bee preference for flowers arranged on a horizontal plane versus inclined planes’, Funct. Ecol. 22(6), 1027–1032.

Manning, A. (1956), ‘The effect of honey-guides’, Behaviour 9(1), 114–139.

Mathis, A., Mamidanna, P., Cury, K. M., Abe, T., Murthy, V. N., Mathis, M. W. & Bethge, M. (2018), ‘Deeplabcut: markerless pose estimation of user-defined body parts with deep learning’, Nat. Neurosci. 21(9), 1281.

Nepi, M., Grasso, D. A. & Mancuso, S. (2018), ‘Nectar in plantinsect mutualistic relationships: From food reward to partner manipulation’, Front. Plant Sci. 9.

Orbán, L. L. & Plowright, C. M. (2014), ‘Getting to the start line: how bumblebees and honeybees are visually guided towards their first floral contact’, Insectes Soc. 61(4), 325–336.

Pasquaretta, C., Jeanson, R., Pansanel, J., Raine, N. E., Chittka, L. & Lihoreau, M. (2019), ‘A spatial network analysis of resource partitioning between bumblebees foraging on artificial flowers in a flight cage’, Movem. Ecol. 7(1).

Penny, J. H. J. (1983), ‘Nectar guide colour contrast: a possible relationship with pollination strategy’, New. Phytol. 95(4), 707–721.

Reber, T., Baird, E. & Dacke, M. (2016), ‘The final moments of landing in bumblebees, bombus terrestris’, J. Comp. Physiol. A. 202(4), 277–285.

Rusch, C., Alberto, D. A. S. & Riffell, J. A. (2021), ‘Visuo-motor feedback modulates neural activities in the medulla of the honeybee, apis mellifera’, J. Neurosci. 41(14), 3192–3203.

Sprengel, C. K. (1793), Das entdeckte Geheimnis der Natur im Bau und in der Befruchtung der Blumen., Friedrich Vieweg, Berlin, Germany.

Srinivasan, M. V., Zhang, S. W., Chahl, J. S., Barth, E. & Venkatesh, S. (2000), ‘How honeybees make grazing landings on flat surfaces.’, Biol. Cybern. 83(3), 171–183.

Stöckl, A. L. & Kelber, A. (2019), ‘Fuelling on the wing: sensory ecology of hawkmoth foraging.’, J. Comp. Physiol. A. 205, 399–413.

Taylor, G. J., Tichit, P., Schmidt, M. D., Bodey, A. J., Rau, C. & Baird, E. (2019), ‘Bumblebee visual allometry results in locally improved resolution and globally improved sensitivity.’, eLife 8, e40613.

Tedore, C. & Nilsson, D.-E. (2019), ‘Avian uv vision enhances leaf surface contrasts in forest environments’, Nat. Commun. 10(1), 238.

Thakar, J. D., Kunte, K., Chauhan, A. K., Watve, A. V. & Watve, M. G. (2003), ‘Nectarless flowers: ecological correlates and evolutionary stability’, Oecologia 136(4), 565–570.

Valenta, K., Nevo, O., Martel, C. & Chapman, C. A. (2016), ‘Plant attractants: integrating insights from pollination and seed dispersal ecology’, Evol Ecol 31(2), 249–267.

van der Kooi, C. J., Dyer, A. G., Kevan, P. G. & Lunau, K. (2019), ‘Functional significance of the optical properties of flowers for visual signalling.’, Ann. Bot. 123, 263–276.

Vorobyev, M., Gumbert, A., Kunze, J., Giurfa, M. & Menzel, R. (1997), ‘Flowers through insect eyes’, Israel J. Plant Sci. 45(2-3), 93–101.

Waser, N. M. & Price, M. V. (1985), ‘The effect of nectar guides on pollinator preference: experimental studies with a montane herb’, Oecologia 67(1), 121–126.

Wolf, S., Roper, M. & Chittka, L. (2015), ‘Bumblebees utilize floral cues differently on vertically and horizontally arranged flowers’, Behav Ecol 26(3), 773–781.

